# The gamma-butyrolactone receptors ScbR and AtrA form a quorum sensing switch between coelimycin and actinorhodin synthesis in *Streptomyces coelicolor* A3(2)

**DOI:** 10.1101/2024.03.11.584493

**Authors:** Bartosz Bednarz, Magdalena Kotowska, Mateusz Wenecki, Marta Derkacz, Adrianna Jastrzemska, Jarosław Ciekot, Krzysztof J. Pawlik

## Abstract

**Background:** Quorum sensing enables gene expression regulation in response to changes in cell population density and controls diverse processes, such as biofilm formation, virulence and antibiotic production, in bacteria. In one of the largest, soil-dominant phylum *Actinobacteria*, cell-to-cell communication occurs through the small, membrane-diffusible signalling molecules gamma-butyrolactones (GBLs). Their actions are exerted through receptor proteins that also act as response regulators in a one-component system manner. With only a few GBL systems characterized, most of them come from the large, antibiotic-producer genus *Streptomyces*. In the model organism *Streptomyces coelicolor* A3(2), two GBL receptors, ScbR and SlbR, which are both antibiotic production repressors, have been reported so far.

**Results:** In this work, we identified a new GBL receptor protein, the conserved and pleiotropic regulator AtrA, which has an activating mode of action. Moreover, we elucidated the precise mechanism by which it controls the production of the antibiotic actinorhodin through the actinorhodin biosynthetic gene cluster activator ActII-orf4. GBL binding to AtrA prevents its binding to the promoter of the *actII-orf4* gene, thereby disabling its transcription, while at the same time, GBL binding to ScbR causes coelimycin antibiotic synthesis derepression.

**Conclusions:** The opposite modes of action of ScbR (repressor) and AtrA (activator) have opposite effects upon GBL binding, activating coelimycin and blocking actinorhodin production at the same time. This phenomenon constitutes an elegant regulatory mechanism that ensures that coelimycin and actinorhodin production are mutually exclusive. These findings also suggest that quorum sensing must be taken into account when designing efficient antibiotic production processes and can be manipulated to ensure both better yield and specificity.

## INTRODUCTION

As one of the dominant soil microbes, *Actinobacteria* play a key role in biogeochemical cycling of carbon, nitrogen, phosphorus and other elements. They produce extracellular hydrolytic enzymes that degrade organic compounds of both plant and animal origin and are essential for soil formation (1). Moreover, members of this phylum are of special relevance to agriculture and the pharmaceutical industry as biocontrol and plant growth-promoting agents (2) as well as as a source of bioactive compounds, antibiotics and enzymes (3). *Streptomyces*, the largest genus of *Actinobacteria*, are mainly soil- and decaying vegetation-dwelling filamentous bacteria that are renowned for producing two-thirds of clinically relevant antibiotics as well as anthelmintic, anticancer and immunosuppressant secondary metabolites. Their complex life cycle involves vegetative growth, aerial mycelium formation and sporulation, processes usually associated with filamentous fungi (3). Because of their diverse ecological niches as saprophytes and the need to integrate and adapt to multiple signals from the environment, *Streptomyces*, which possess one of the most complex regulatory machineries in the bacterial kingdom, have become an outstanding model for genome regulation studies.

The synchronized behavior of bacteria is regulated by a cell-to-cell signalling system called quorum sensing, which controls bacterial swarming, virulence factor expression, exopolysaccharide biosynthesis (4), morphogenesis and secondary metabolite production (5). Quorum-sensing systems consist of membrane-diffusible molecules and their receptor proteins, which also act as response regulators. N-acylhomoserine lactones (AHLs) are the prevalent quorum sensing molecules in gram-negative bacteria. AHL receptors are LuxR family proteins that typically activate transcription by binding to promoters of their target genes when in complex with signalling molecules. Gram-positive bacteria use processed oligopeptides (autoinducing polypeptides) or gamma-butyrolactones (GBLs) and related classes of butenolides and furans for cell-to-cell communication (4–6). GBL receptors generally belong to the TetR family of regulators, typically repressors, which dissociate from DNA upon GBL binding. A recent study by Liu and coworkers revealed the widespread response of AHL receptors from gram-negative bacteria to GBLs from *Streptomyces* (7).

The *S. coelicolor* A3(2) chromosome encodes enzymes for the production of at least 20 secondary metabolites, four of which have antimicrobial activities: coelimycin A (CPK A), actinorhodin (ACT), undecylprodigiosin (RED) and the calcium-dependent antibiotic (CDA) (8). The production of these compounds must be tightly controlled, taking into account the timing (cell density and differentiation state), nutrient availability and competition or symbiosis with other organisms. Therefore, many levels of regulation exist, which are exerted in a cascade manner through the so-called “pleiotropic” and “pathway-specific” regulators (9). The most immediate regulators, *Streptomyces* antibiotic regulatory proteins (SARPs), directly activate the transcription of biosynthetic enzymes, and their genes are colocalized with those of biosynthetic enzymes on the chromosome in biosynthetic gene clusters (BGCs). The pathway-specific activators of coelimycin synthesis (*cpk* cluster) are CpkO and CpkN, calcium-dependent antibiotic (*cda* cluster) - CdaR, undecylprodigiosin (*red* cluster) - RedZ and RedD, and actinorhodin (*act* cluster) - ActII-orf4 (9). The transcription of SARP genes is controlled by upper-level regulators belonging to different families. One such protein, the TetR-like pleiotropic regulator ScbR, is a well-studied gamma-butyrolactone receptor that is a direct repressor of the coelimycin synthesis activator gene *cpkO* (10,11). When the cell density and GBL level reach the threshold at approximately 20–24 hours of *S. coelicolor* A3(2) culture growth, ScbR dissociates from the *cpkO* promoter upon GBL binding, which results in coelimycin production activation (12–14). The precise mechanism of this process was recently reviewed by our group (15). The second GBL receptor, SlbR, which does not belong to the TetR family, was identified in this organism, although no specific regulatory mechanism was identified (16–18). Two other proteins, the pleiotropic TetR-like regulators CprA and CprB, were predicted to bind gamma-butyrolactones in *Streptomyces coelicolor* A3(2) based on their high amino acid sequence identity to the *Streptomyces griseus* GBL receptor ArpA; however, there is no experimental evidence to date. The proteins exhibit somewhat contrary activities: CprA acts as an activator, while CprB downregulates antibiotic synthesis in this strain (19) This work identified a novel gamma-butyrolactone receptor in *S. coelicolor* A3(2), AtrA, which until now has been a well-known pleiotropic regulator and activator of the actinorhodin biosynthetic gene cluster. Previously, *atrA* deletion was characterized by a near-complete reduction of actinorhodin production, and this effect was reversed in the *atrA* complementation strain of *S. coelicolor* A3(2). This phenotype was accompanied by corresponding changes in *actII-orf4* transcription, as detected by qPCR. The exact sequences of the two AtrA binding sites in the *actII-orf4* promoter/coding sequence were determined by footprinting. It was also noted that *S. coelicolor* A3(2) AtrA was able to bind to the promoter sequence of the *S. griseus strR* gene, encoding the final regulator of streptomycin production in this distant relative strain (20). Later, it was confirmed that *S. coelicolor* A3(2) AtrA is capable of influencing streptomycin production in *S. griseus*, but surprisingly, the effect was inhibitory. The authors noted that in *S. coelicolor* A3(2), AtrA might have coevolved with different target sequences than those present in *S. griseus* and hence is unable to sufficiently activate the *strR* promoter or even obstructs the binding of native *strR* activators in the latter strain (21). Hirano et al. further tested the binding of AtrA to *strR* but used the *S. griseus* protein. Indeed, the interaction was observed, but AtrA was not essential for streptomycin production. However, it binds to the *strR* promoter between two sites of the essential, pleiotropic activator AdpA and enhances streptomycin biosynthesis (22).

AtrA was also found to directly activate transcription of the *nagE2* permease gene in *S. coelicolor* A3(2) (23). NagE2 is the transporter of N-acetylglucosamine as part of the phosphotransferase system (PTS), which is involved in carbon source uptake and carbon signalling in bacteria. N-acetylglucosamine signalling, an indication of cell wall peptidoglycan autolysis, is crucial for the development and antibiotic production of *S. coelicolor* A3(2). The uptake of this chemical halts both sporulation and antibiotic synthesis on nutrient-rich media while promoting the same processes under poor nutritional conditions (24,25). Interestingly, *nagE2* was shown to peak in transcription at approximately the same time as *actII-orf4,* and the transcription of both genes was strongly reduced in the *atrA* deletion strain, providing clear evidence that this gene is a part of the AtrA regulon (23).

Later, it was found that AtrA could bind to the intergenic region between *atrA* and *SCO4119* at a site that suggested the activation of *SCO4119*, which was confirmed at the transcriptional level. However, the protein also bound to the promoter sequence near its own gene, as well as in the middle of the intergenic region, which left the possibility of autoregulation unverified (26).

In *S. coelicolor* A3(2), SsgR is the activator of *ssgA,* encoding the protein responsible for controlling the localization of the SsgB protein, which recruits FtsZ to septation sites during sporulation (27,28). In this species, the *ssgR* gene was shown to be directly activated by AtrA, which was subsequently corroborated by the downregulation of the *ssgR* transcript in the *atrA* null strain (29).

Similarly, the AtrA homologue from *Streptomyces avermitilis*, AveI, positively regulates morphological differentiation by activating *ssgR* and *ssgD* and repressing *wblI* (30). AveI was shown to directly regulate 35 genes. It promoted melanin synthesis by activating the *melC1C2* operon but inhibited oligomycin production by repressing the cluster-situated regulator *olmRII* and the structural gene *olmC.* It also repressed the production of avermectin by binding to the promoters of the cluster-situated regulator *aveR* and the structural genes *aveA1*, *aveA3* and *aveD*. Notably, in an early study of AveI, its binding to AveR could not be detected (30). It was shown, however, that AveI can replace AtrA in *S. coelicolor* A3(2) and activate actinorhodin synthesis and that AtrA functions in *S. avermitilis* in the same way as AveI in downregulating avermectin production (31)

The AtrA homologue from *Streptomyces pristinaespiralis* was demonstrated to positively influence pristinamycin I and II synthesis in this species by directly activating the cluster-situated regulatory genes *spbR* (a gamma-butyrolactone receptor) and *papR5* (encoding a TetR-like repressor) (32).

In a recent study, the positive effect of the AtrA homologue on lincomycin production was characterized in *Streptomyces lincolnensis*. Disruption of the gene downregulated the transcription of the cluster-situated regulator *lmbU* and the structural genes *lmbA* and *lmbW*. It also resulted in the downregulation of *adpA* and *bldA* transcription. We also direct readers to this publication for amino acid sequence comparison of the AtrA homologues that are discussed in this introduction (33).

Mao and colleagues demonstrated that the AtrA homologue in *Streptomyces roseosporus* activated the production of daptomycin by directly binding to the *dptE* promoter. The *atrA* gene promoter was itself under the direct positive control of the AdpA homologue, which in turn was directly repressed by the homologue of *S. griseus* GBL-binding ArpA. The group showed that in this strain, deletion of *atrA* and deletion of *adpA* both result in a bald phenotype and loss of daptomycin production. On the other hand, deletion of *arpA* accelerated the production of daptomycin in *S. roseosporus*. Taken together, these findings demonstrate that in *S. roseosporus*, AtrA is part of a GBL signalling pathway in which GBLs induce ArpA dissociation from the *adpA* gene promoter, resulting in its derepression. AdpA then activates *atrA,* which in turn activates daptomycin synthesis by binding to the *dptE* promoter. AtrA autoactivation of *atrA* transcription serves as a positive feedback loop in this pathway (34). Recently, *S. roseosporus* AtrA was shown to be recognized by ClpX and directed to the ClpP protease for degradation. The authors also mutated the recognized AAA motifs of AtrA to increase the yield of daptomycin in the strain (35).

Importantly, in *Streptomyces globisporus*, the AtrA homologue serves as part of a negative feedback loop in which the protein directly activates the transcription of the lidamycin biosynthesis-related CSR gene, *sgcR1,* and this interaction is in turn inhibited by AtrA binding to heptaene, a lidamycin production intermediate. Interestingly, in the same paper, the protein was shown to bind *S. coelicolor* A3(2) actinorhodin, which caused its dissociation from the promoter DNA. However, which form of actinorhodin could be the actual ligand has not been determined (36). Later, Al-Tarawni (37) concluded that γ-actinorhodin is a nonspecific inhibitor of multiple transcription factors, including AtrA from *S. coelicolor* A3(2).

In his effort to assess the ability of actinorhodin to interact with *S. coelicolor* AtrA, Hassan (38) unexpectedly observed that a crude extract from the M1146 strain (*Δact*, *Δred*, *Δcpk*, and *Δcda* quadruple mutant), which was used as a negative control, inhibited the DNA binding activity of AtrA. However, the active compound was not identified. Here, we demonstrate that the mysterious molecule(s) that interact with AtrA are γ-butyrolactone(s), which are signal transduction molecule(s).

## MATERIALS AND METHODS

### Bacterial strains and growth conditions

The bacterial strains used in this study are listed in Supplementary Table S1. For genetic manipulation, *Escherichia coli* and *Streptomyces coelicolor* A3(2) strains were grown in standard conditions (39,40). For visual imaging of the strain phenotypes (spots), 20 μl of spore suspension in water (OD_600_=0.3) was spotted on solid medium 79 without glucose (79NG) (41) and grown at 30°C. To demonstrate the activity of exogenous GBLs, the *S. coelicolor* double mutant Δ*scbA-atrA_OE_* was grown as a confluent lawn on solid 79NG medium for 18 h, after which 10 µl of extract from a 48-h-old culture of either Δ*cpkO* (GBL-containing extract) or Δ*scbA* (negative control) was added to the center of the plate, incubated further and photographed 4, 7 and 23 h after GBL addition. For RNA isolation and actinorhodin production studies, 200 μl of spore suspension in water (OD_600_=0.3) was spread on solid 79NG medium overlaid with perforated cellophane and grown at 30°C. Biomass scraped from half of the surface of the cellophane from each plate and frozen in liquid nitrogen was used for RNA isolation and qPCR analysis. For liquid cultures, 50 ml of Glu-MM medium (42) in 250 ml Erlenmeyer flasks with springs was inoculated with spores to an OD_600_ of 0.1 and incubated at 30°C on an orbital shaker (220 rpm).

### *S. coelicolor* A3(2) *atrA* deletion and overexpression mutants

The constructs and oligonucleotides used in this study are listed in Supplementary Tables S2 and S3, respectively. PCR-amplified DNA fragments were either inserted into the appropriate plasmids using the Gibson assembly technique (43) or first cloned into the pUC18 vector (SmaI sites) and then moved into the appropriate vector by restriction cloning. All of the introduced sequences were verified by sequencing. The constructs were subsequently introduced into *S. coelicolor* A3(2) via *E. coli* ET12567/pUZ8002-mediated conjugation (39) For the *atrA* deletion construct StD72A-atrA_DM_, the *atrA* gene sequence in the cosmid StD72A was substituted using PCR targeting (44) with the *aac3(IV)* apramycin resistance cassette. The modified cosmid was subsequently introduced into *S. coelicolor* M145. Transconjugants were screened for apramycin resistance and kanamycin sensitivity to obtain the *atrA* deletion strain *ΔatrA* (P332). The pIJ10257-atrA_OE_ construct containing the *atrA* gene under the control of the *ermEp\** promoter was introduced into the wild-type (M145) and *ΔscbA* (M751) (11) strains to obtain the M145-*atrA*_OE_ (P330) and *ΔscbA*-*atrA*_OE_ (P331) strains, respectively. The empty vector pIJ10357 was introduced into the M145, *ΔscbA* and *ΔatrA* strains to obtain the control strains M145-Φ (P132), *ΔscbA*-Φ (P053), and *ΔatrA*-Φ (P334), respectively.

### Measurement of actinorhodin production

The actinorhodin assay was adapted from Kieser et al. (39). Whole agar medium (20 ml) from a Petri plate, from which the cellophane disc was removed, was transferred to a 50 ml Falcon tube, and 10 ml of a methanol-chloroform (vol. 1:1) mixture was added. The sample was extracted for 30 min on a roller at room temperature and centrifuged (10 min, 4800 × g). One hundred microliters of the upper phase was mixed with 5 µl of 5 M KOH. The absorbance at 640 nm was measured with a ClarioStar Plus microplate reader (BMG Labtech).

### RNA isolation, reverse transcription and quantitative PCR

*S. coelicolor* A3(2) RNA was isolated using a GeneJet RNA Purification Kit (Thermo) with a modification in the biomass disruption step. On ice, in a 2 ml Eppendorf tube, the frozen biomass was suspended by pipetting in 100 µl of buffer TE with 15 mg/ml lysozyme. Then, 300 µl of lysis buffer supplemented with DTT (as described in the manual) was added. Next, approximately 200 µl of 1 mm-diameter glass beads was added, and the sample was homogenized using a BeadBug6 homogenizer (Benchmark) in one 30 s cycle at 4000 rpm. The samples were centrifuged at 12000 × g for 1 min, applied to purification columns and processed according to the manufacturer’s instructions. Elution was performed in 50 µl of nuclease-free H_2_O. The removal of genomic DNA was performed by subsequent on-column purification according to the manufacturer’s instructions. RNA integrity was assessed by applying ∼200 ng of total RNA to a 0.5× TBE agarose gel (1%). Prior to electrophoresis, the samples were mixed with the loading dye and incubated at 70°C for 5 min.

Reverse transcription was performed with a Maxima First Strand cDNA Synthesis Kit for RT-qPCR (Thermo Fisher Scientific) using random hexamers. The total volume of the reaction was 10 μl, with the amount of DNase-treated RNA ranging from 200 ng to 800 ng. Reaction conditions were modified for a GC-rich RNA as follows: 10 min at 25°C, 20 min at 65°C, and 5 min at 85°C. After the reaction, depending on the amount of RNA used, samples with less than 270 ng of template RNA were diluted with miliQ water 12 times (120 μl final), whereas samples with more than 270 ng were diluted 20 times (200 μl final). cDNA was stored at −80°C. In qPCR, Cq values for *hrdB* transcripts in samples after reverse transcription (RT+) and in control samples lacking the enzyme in reverse transcription reaction (RT-) were compared for assessment of genomic DNA contamination. For qPCR, RNA/cDNA samples with a Cq above 32 for RT-reactions and at least 5 cycles difference between RT+ and RT-reactions were used.

qPCR was performed on a Bio-Rad CFX96 apparatus in Hard-Shell® 96-well PCR low-profile plates (Bio-Rad, cat. no: HSP9601) with Optical Microseal ‘B’ PCR Plate Sealing Film (Bio-Rad, cat. no: MSB1001). PowerUp SYBR Green Master Mix (Applied Biosystems) was used for the reactions. The assay was performed for 3 biological replicates with at least 3 technical replicates for each biological replicate. For reactions with a total volume of 15 μl, 4 μl of cDNA per reaction was used. The program settings were as follows: 2 min at 50°C; 2 min at 95°C; 40 cycles of 15 s at 95°C, 30 s at 60°C, and 30 s at 72°C. After each PCR, melting curve analysis was performed. The primer sets for the targeted genes were validated by creating a standard curve with 10-fold dilutions of template material. In the reactions with 0.5 μM final primer conc. and an annealing temperature of 60°C, we achieved R^2 = 0.99 and efficiencies of 97.71% for *hrdB*, 101.56% for *atrA*, 93.69% for *actII-orf4* and 91.69% for *cpkO*. The relative gene expression ratio to that of *hrdB, which was used* as an endogenous control, was calculated using the Pfaffl formula (45).

### Purification of recombinant proteins

Purification of His-tagged ScbR and ScbR2 proteins was performed as described previously (12). Glutathione S-transferase (GST), expressed from the pGEX-6P-2 plasmid in *E. coli* BL21 and purified under standard conditions (GST Gene Fusion System Handbook 18-1157-58, Amersham Biosciences), was a kind gift from Jakub Muraszko.

For His-tagged AtrA purification, 1 L of LB medium supplemented with kanamycin (50 μg/ml) was inoculated with 10 ml of an overnight culture of *E. coli* BL21(DE3)Star with the pET28a-atrA plasmid and shaken (180 rpm) at 37°C until the OD_600_ reached 0.65. The expression was induced with 1 mM IPTG, and incubation was continued for 3 h. The biomass was centrifuged (5000 × g for 10 min) and frozen at −20°C. The cells were resuspended in 20 ml of lysis buffer (50 mM Tris-HCl pH 8, 300 mM NaCl, 1 mM DTT, 0.01% Tween 20, 20 mM imidazole) supplemented with cOmplete Mini EDTA-Free Protease Inhibitor Cocktail (Sigma) and lysozyme (Fluka) (1 mg/ml) and incubated at room temperature for 15 min. The cells were disrupted at 20 kPsi (one shot) in Cell Disruptor (Constant Systems Ltd.) and centrifuged (20000 × g, 30 min, 4°C). The supernatant was applied to a 1 ml HisTrap FF column (GE Healthcare) by a peristaltic pump. The column was washed with 15 column volumes (CV) of lysis buffer, and then the protein was eluted with lysis buffer supplemented with 50 mM imidazole. The AtrA-containing fractions were pooled, and glycerol was added to a final concentration of 50%. The samples were stored at −20°C.

For His-tagged SlbR purification, two 5 L flasks, each containing 1 L of LB medium with kanamycin (50 μg/ml), were inoculated with 10 ml of an overnight culture of *E. coli* BL21 (DE3)Star containing the pET28a-slbR plasmid and shaken (180 rpm) at 37°C until the OD_600_ reached 0.6. The expression was induced with 1 mM IPTG, and incubation was continued for 6 h at 30°C. Biomass was centrifuged (5000 × g for 10 min) and frozen at −20°C. The cells were resuspended in 80 ml of lysis buffer (50 mM Tris-HCl pH 8, 500 mM NaCl, 1 mM DTT, 0.01% Tween 20, 10% glycerol, 0.01% β-mercaptoethanol, 20 mM imidazole) supplemented with protease inhibitors and lysozyme (1 mg/ml) and incubated at room temperature for 30 min. The cells were disrupted at 20 kPsi (one shot) in a cell disruptor and centrifuged (20000 × g, 30 min, 4°C). One milliliter of equilibrated HIS-Select Nickel Affinity Gel (Sigma) was added to the cleared cell lysate and incubated with mild agitation for 1 h at 4°C. The resin was centrifuged (500 × *g*, 5 min, 4°C), washed three times with lysis buffer and transferred to an empty column. The protein was eluted with lysis buffer supplemented with 100 mM imidazole. SlbR-containing fractions were pooled, and glycerol was added to a final concentration of 50%. The samples were stored at −20°C.

### γ-butyrolactone extraction

GBL extracts were prepared from 450 ml of *S. coelicolor* A3(2) culture supernatant as described previously (46). Cultures grown in liquid Glu-MM medium were centrifuged at 24000 × g for 10 min. The supernatants were mixed with 3 volumes of ethyl acetate (POCH) and shaken 10 times. The upper ethyl acetate phase was collected and evaporated under vacuum. The residue was dissolved in 500 µl of methanol and stored at −20°C. For the EMSAs, 500 µl of the extracts were once again evaporated with a Speedvac and dissolved in 250 µl of methanol. For the GBL capture experiment, GBL-containing extract was evaporated with a Speedvac and dissolved in an equal volume of water.

### GBL affinity capture

The binding of GBLs to recombinant receptor proteins was assessed by the affinity capture method adapted from (47). Equimolar quantities of ScbR, ScbR2, AtrA, SlbR and GST proteins, corresponding to 1.5 mg of ScbR, were washed three times with 3 ml of water using Amicon Ultra-4 (3 kDa cut-off, Millipore) (centrifugation at 4°C, 4800 × g). The concentrated protein sample (approx. 150 μl) was mixed with water (up to 710 μl), 40 μl of 10x LI-COR buffer (100 mM Tris pH 7.5, 500 mM KCl, 10 mM DTT) and 50 μl of GBL solution (extract from the M1154 strain grown for 72 h). After 12 h of gentle agitation at room temperature, the samples were subjected to ultrafiltration on the same Amicon Ultra-4 filters by adding four 3 ml portions of water. The sample volume was adjusted to 800 μl with water, and 2 μl of 6 M HCl was added. The mixtures were denatured at 95°C for 3 min and ultrafiltered. The filtrate was collected, concentrated 40 times with a Speedvac and subjected to HPLC-MS for GBL detection. The control sample without protein was processed in the same way.

### HPLC-MS analysis of γ-butyrolactones

The apparatus used was a Dionex 3000 RS-HPLC instrument equipped with a DGP-3600 pump, a WPS-3000 TLS TRS autosampler, a TCC-3000 RS column compartment (Dionex Corporation, USA) and a micrOTOF-QII mass spectrometer as a detector (Bruker Daltonics, Germany). The chromatography column used was a 50 x 2.1 (i.d.)-millimeter Thermo Scientific Hypersil Gold column with 1.9-micron particles (Part No. 25002-052130, Serial No. 0110796A6). The sample injection volume was 1 µl. The flow rate was 0.2 ml/min, and the eluate was monitored by mass spectrometry. The mobile phase was 0.1% formic acid in water (solvent A) and 0.1% formic acid in acetonitrile (solvent B). The ramp was as follows: 0 min – 5% B, 0.5 min – 5% B, 20.5 min – 95% B, 22.5 min – 95% B, and 22.6 min – 5% B. The mass spectrometer was calibrated with 10 mM sodium formate, and the following settings in positive ESI mode were used. Scan range: 100-1500 m/z, end plate offset: −500 V, capillary voltage: −4000 V, nebulizer gas (N2): 1.2 bar, dry gas (N2): 9 L/min, dry temperature: 180°C. To obtain information about GBL1-GBL8 abundance in the analysed samples, positive ion mode chromatograms were extracted based on the calculated m/z values for the four ions ([M+Na]+, [M+H]+, [M-H2O+H]+, [M-2H2O+H]+) of GBLs (SCB1-SCB8), and peak areas were calculated. GBLs with identical molecular masses were distinguished on the basis of their relative retention times according to (48).

### Electrophoretic mobility shift assay (EMSA)

The promoter region of *actII-orf4* was amplified with the primers pTZXBA700 and pTZBAM700 from the pTZ-pactII-orf4 plasmid as a template. The fragment encompassing the promoters of both the *atrA* and *SCO4119* genes was amplified with the primers EMSAFLUXF-FAM and EMSAFLUXR-FAM, and the pFLUXH-patrA plasmid was used as a template. Each EMSA sample contained either 0.03 pmol of the IRDye 700-labelled pactII-orf4 fragment or 0.06 pmol of the FAM-labelled patrA/SCO4119 fragment and variable amounts of AtrA protein and GBL containing/control extracts, as indicated. Other buffer components and conditions were as described previously (12). Gels were visualized using a Typhoon FLA9500 (GE Healthcare).

## RESULTS

### Abundant production of GBLs by the Δ*cpkO* and M1154 strains

One of the effects of *cpkO* gene deletion, encoding the main SARP activator of the coelimycin biosynthetic gene cluster, is an over 20-fold increase in the abundance of the GBL synthase ScbA resulting from the absence of the ScbR2 repressor, which normally shuts down its expression (12). Increased GBL production was observed in the antibiotic production superhost *S. coelicolor* M1152 (49), from which four BGCs were deleted, including the *cpk* cluster with the *scbR2* gene, while *scbA* (and the accessory gene *scbB*) were left behind (48). In the present study, we used both the Δ*cpkO* and M1154 strains as sources of GBLs. Strains M1152 and M1154 are both quadruple BGC deletion mutants derived from M1146 (*Δact*, *Δred*, *Δcpk*, *Δcda*) by introducing one or two point mutations in the *rpoB* and *rpsL* genes (49).

Here, we developed a simple biological assay for GBL production in liquid cultures. The Δ*scbA* strain, which is unable to synthesize GBLs, served as a reporter. A lack of GBLs provides complete repression of the *cpk* gene cluster by the ScbR protein. The addition of external GBLs leads to ScbR dissociation from DNA and allows coelimycin production. The use of Glu-MM medium ensures that all CPK precursors turn into the yellow glutamate adduct coelimycin P2 (42). Twenty-two-hour cultures of Δ*scbA* were centrifuged, and the biomass was mixed with spent culture media from Δ*cpkO* and M1154, which are not able to produce CPK. Medium from Δ*scbA* culture was used as a negative control. After several hours of additional cultivation, yellow pigment production was visible in the cultures in which GBLs were present (Fig. 1). As expected, media from both Δ*cpkO* and M1154 cultures derepressed CPK production in Δ*scbA*. An obvious limitation of the assay is the high concentration of actinorhodin in the tested culture media. However, the slightly blue color of the medium from the 48 h culture of Δ*cpkO* did not prevent the observation of CPK.

**Fig. 1.**
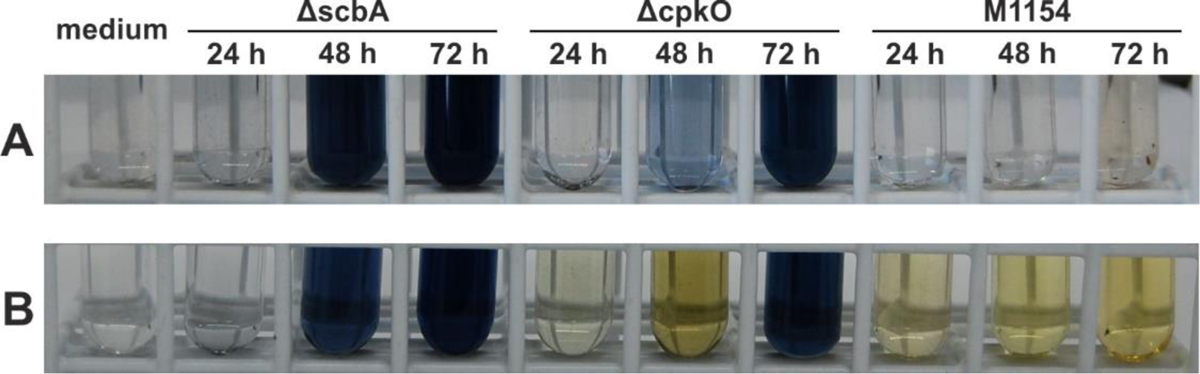
GBLs present in the *ΔcpkO* and M1154 culture media induce coelimycin (yellow pigment) production by the *ΔscbA* strain. (A) Supernatants of *ΔscbA, ΔcpkO* and M1154 cultures in Glu-MM medium collected after the indicated time of growth. (B) The same supernatants after mixing with *ΔscbA* biomass and further incubation for 8 h.

GBL-containing extracts from the culture media of Δ*cpkO* and M1154 were also analysed by HPLC-MS. The Δ*scbA* strain was used as a negative control. The presence of all gamma-butyrolactones (SCB1-8) reported by Sidda et al. (48) was confirmed in extracts from 48 h and 72 h cultures of both strains (Fig. S1). Interestingly, three other putative GBLs, tentatively named SCB9, SCB10 and SCB11, were also detected. The observed molecular mass of compound SCB9 was identical to that of SCB1 and SCB2, while the mass of compounds SCB10 and SCB11 was the same as that of SCB3 and SCB7.

### AtrA is a gamma-butyrolactone receptor

To determine whether AtrA could be a GBL receptor, we performed an electrophoretic mobility shift assay (EMSA) with the protein and its two target DNA fragments reported earlier, the *actII-orf4* gene promoter (20) and the intergenic region between *atrA* and its neighbor *SCO4119* (26) (Fig. 2). The addition of GBL-rich extract prevented the binding of both DNA fragments. This effect could not be observed when the same volumes of extract from the Δ*scbA* strain (lacking GBLs), extract from sterile Glu-MM medium or methanol were used. These results indicate that AtrA is a gamma-butyrolactone receptor that dissociates from its target sequence upon GBL binding.

**Fig. 2.**
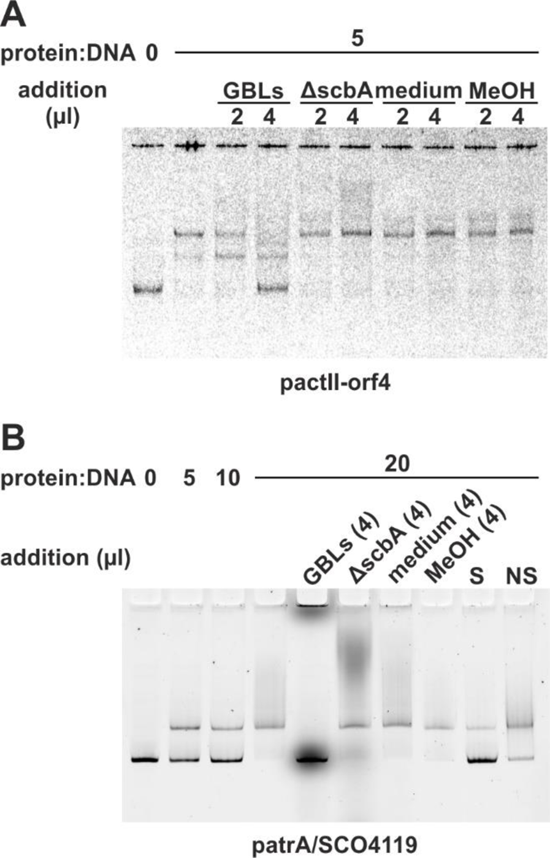
Electrophoretic mobility shift assay. AtrA binding to the *actII-orf4* (A) and *atrA/SCO4119* (B) promoter regions and its dissociation upon the addition of γ-butyrolactones (GBLs). Extracts from the ΔscbA strain and Glu-MM medium, as well as methanol (MeOH), were used as controls. S, NS – 10-fold excess of specific and nonspecific competitor DNA, respectively.

To directly assess the ability of AtrA to bind GBLs, an affinity capture method adapted from (47) was applied. Extract from the M1154 strain grown for 72 h was used as a source of GBLs. It contained a mixture of nine GBLs (Fig. S2 and 3A). Two known GBL receptors from *S. coelicolor* A3(2), ScbR and SlbR, were used as positive controls. ScbR2, known as a “pseudo” GBL receptor, and the GST protein, which is not associated with the GBL system, were chosen as negative controls. As shown in Fig. 3B and C (GBL capture), the amounts of GBLs detected in the AtrA-bound sample exceeded those of SlbR, a positive control. This confirms that AtrA is indeed able to bind gamma-butyrolactones. As expected, ScbR exhibited the greatest ability to bind different GBLs. Unexpectedly, ScbR2 was also able to bind GBLs. Four compounds, namely, SCB6, 8, 9 and 10, were not detected in any of the protein-bound samples.

**Fig. 3.**
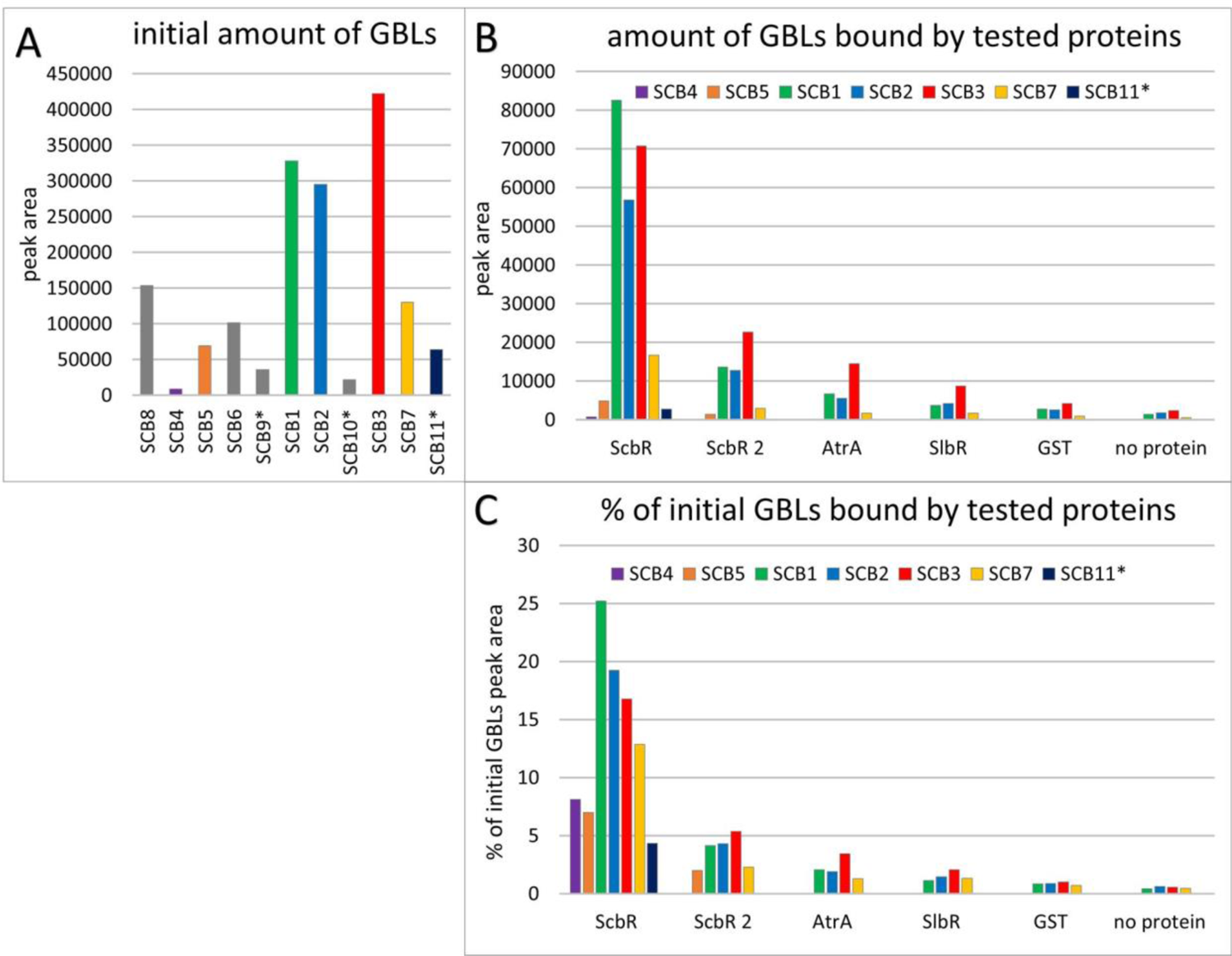
Affinity capture of GBLs by recombinant proteins. (A) GBLs detected by HPLC-MS in the extract from M1154 culture, quantified as peak areas from extracted ion chromatograms. The compounds were ordered according to their retention times. The putative GBLs reported for the first time in the current work are marked with asterisks. The gray columns denote GBLs not detected in the protein-bound samples. (B) GBLs captured by the recombinant proteins incubated with the GBL-containing extract characterized in panel A quantified as peak areas from the extracted ion chromatograms. (C) GBLs captured by the recombinant proteins; peak areas from panel B are expressed as percentages of the respective peak areas from panel A.

### Actinorhodin production is dependent on both AtrA and quorum sensing

#### ΔatrA

Our deletion mutant Δ*atrA* exhibited a complete loss of ACT production when it was grown directly as a spot or on perforated cellophane on solid medium 79NG (Fig. 4 and 5A, B), indicating that AtrA is required for the cascade of actinorhodin synthesis activation. qPCR revealed that the *actII-orf4* transcription of this strain decreased, with fold changes (FCs) of 0.25 and 0.51 in comparison to that of M145 at 27 h and 46 h, respectively (Fig. 5C).

**Fig. 4.**
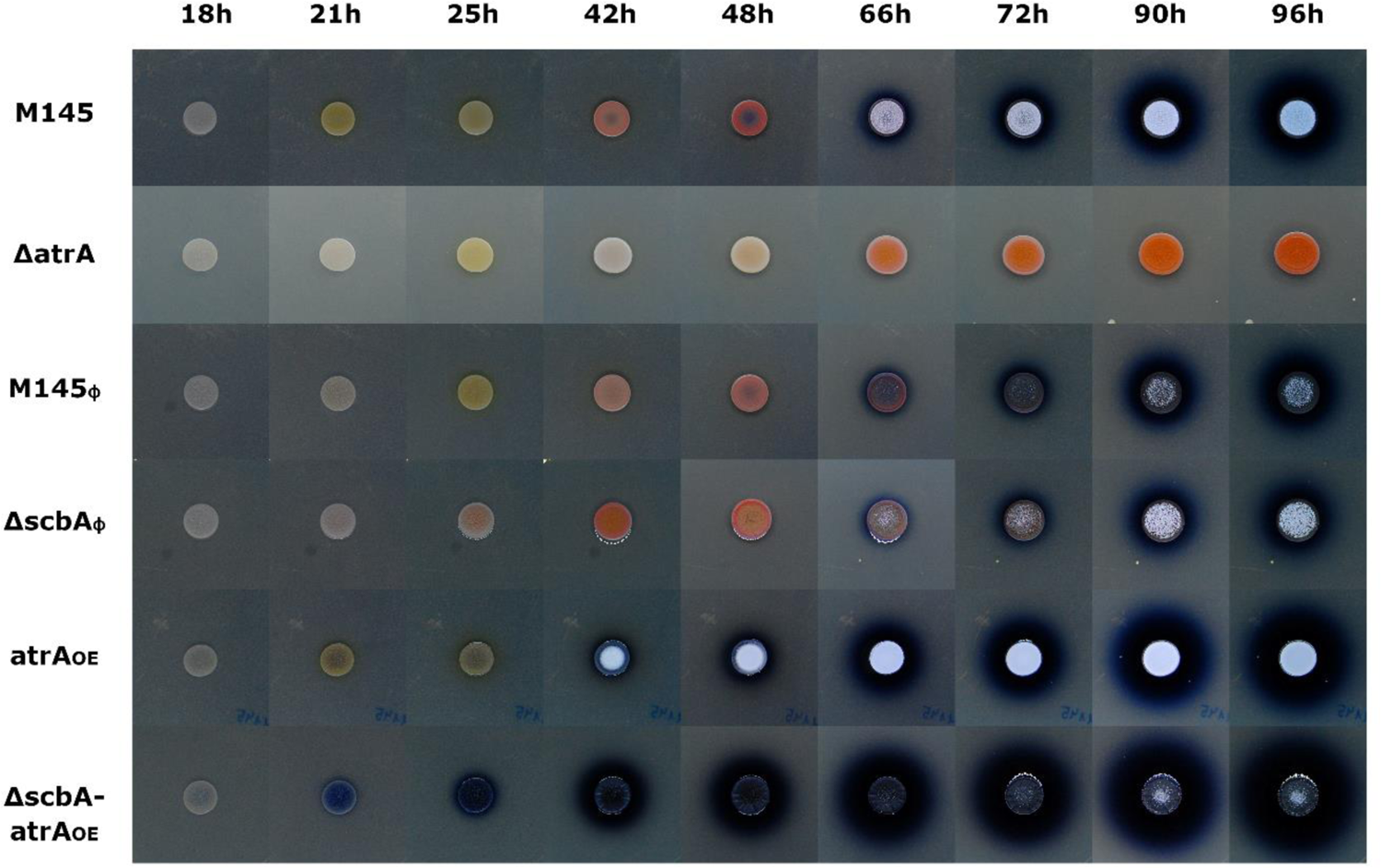
Phenotypes of the *S. coelicolor* A3(2) strains grown on solid medium 79NG as spots.

**Fig. 5.**
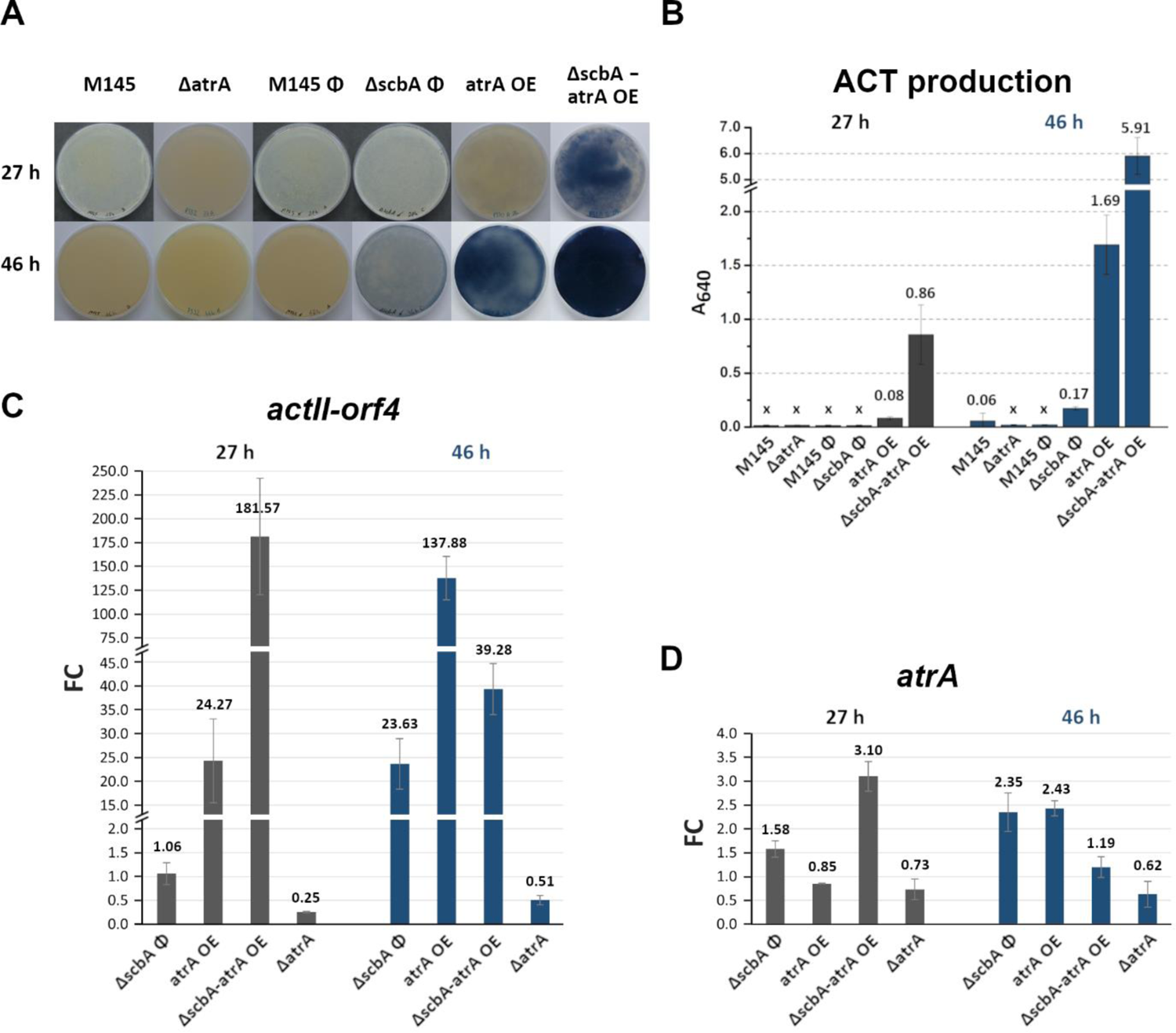
Phenotypes (A), actinorhodin yield (B) and RT-qPCR transcriptional analysis of the *actII-orf4* (C) and *atrA* (D) genes in *S. coelicolor* A3(2) mutant strains grown on 79NG medium covered with cellophane disks. “X” in panel B denotes no visibly detectable production of ACT. For the *atrA* gene (panel D), qPCR primers were designed to specifically detect only the native transcript (from the *atrA* promoter) and not that resulting from overexpression. Gene expression fold changes for strains in panels C and D were calculated as comparisons against their respective control strains: M145 for Δ*atrA* and M145φ for *atrAOE,* Δ*scbA*φ and Δ*scbA*-*atrA*OE.

Although Δ*atrA* was delayed in RED production, it synthesized plenty of this pigment at later timepoints when the strains were grown as spots (Fig. 4).

#### *atrA*_OE_

To further test the activity of AtrA, we generated the *atrA* overexpression mutant *atrA_OE_* and observed that this strain produced ACT much earlier than M145φ, with the blue color filtering through yellow coelimycin as early as at 21 and 25 h, when the strains were grown as spots (Fig. 4). *atrA*_OE_ was also a very potent ACT producer on cellophane disks. ACT synthesis was already visible after 27 h (Fig. 5A) with A_640_=0.08 (Fig. 5B). A very high rate of ACT production was even more evident after 46 h (A_640_=1.69). The transcription of *actII-orf4* matched these phenotypes, with FC values of 24.27 and 137.88 at the 27 and 46 h timepoints, respectively, in comparison to M145φ, which did not yet produce visible quantities of ACT (Fig. 5C).

#### ΔscbA-atrA_OE_

If GBLs indeed bind to AtrA and inhibit its activity, the effect of *actII-orf4* transcription activation by AtrA should be reinforced in a double mutant with the deletion of GBL synthase *scbA* in addition to *atrA* overexpression. This effect became evident as early as at the 18 h timepoint when the Δ*scbA*-*atrA*_OE_ spot culture started to precociously produce actinorhodin and continued throughout the 96 h of growth to accumulate the highest levels of ACT among the studied strains (Fig. 4). Δ*scbA*-*atrA*_OE_ was also the most potent ACT producer on cellophane disks, with enormous yields of A_640_=0.86 and 5.91 at 27 h and 46 h, respectively (Fig. 5A and B). At the transcriptional level, at the 27 h timepoint, Δ*scbA*-*atrA*_OE_ double mutant’s fold change of *actII-orf4* transcript (FC=181.57 relative to M145φ) was much higher than that of the *atrA*_OE_ strain with *atrA* overexpression alone (FC=24.27 relative to M145φ) (Fig. 5C). Interestingly, at the 46 h timepoint, the increase in *actII-orf4* transcription in the Δ*scbA*-*atrA*_OE_ strain (FC=39.28) was not as strong as that in the *atrA*_OE_ strain (FC=137.88), despite the greatest increase in ACT yield (near-black color of agar plates) in the double mutant (Fig. 5A, B, C). We suspect that the accumulation of ACT at such a high concentration may interfere with DNA binding by AtrA.

#### ΔscbAφ

According to the proposed mechanism, even deletion of the GBL synthase *scbA* alone should increase *actII-orf4* transcription by allowing *atrA* to uninterruptedly bind (and activate) its target promoter. This effect could be observed on cellophane discs at the 46 h timepoint, when ACT production was greater in ΔscbAφ (A_640_=0.17) than in M145φ (ACT not detectable) (Fig. 5A and B). The *actII-orf4* transcription of the strain was 23.63 times greater than that of M145φ; however, no significant difference was detected at the 27 h timepoint for either the ACT production phenotype or the *actII-orf4* transcript (Fig. 5A, B, C). This effect was also not observed when bacteria were grown as spots, in which case ACT was produced somewhat faster in M145φ than in Δ*scbA*φ (Fig. 4).

### Gamma-butyrolactones coordinate coelimycin and actinorhodin synthesis

To confirm that gamma-butyrolactones inhibit AtrA-mediated activation of actinorhodin production, we grew the Δ*scbA-atrA*_OE_ double mutant as a confluent lawn on solid 79NG medium for 18 h, after which 10 µl of GBL extract from either the Δ*cpkO* or the Δ*scbA* strain was added to the center of the plate, incubated further and photographed 4, 7 and 23 h after GBL addition. The Δ*cpkO* extract is rich in GBLs because of strongly upregulated *scbA* transcription in this strain, while the Δ*scbA* extract contains no GBLs because of GBL synthase gene deletion (12). The Δ*cpkO* GBL extract prevented ACT production and instead promoted CPK synthesis as seen after 25 and 41 h of Δ*scbA-atrA*_OE_ growth, while the Δ*scbA* extract produced no distinct phenotype (Fig. 6A). Fig. 6B shows a schematic representation of GBL-mediated regulation of coelimycin and actinorhodin synthesis through the action of the TetR-like GBL receptors ScbR and AtrA.

**Fig. 6.**
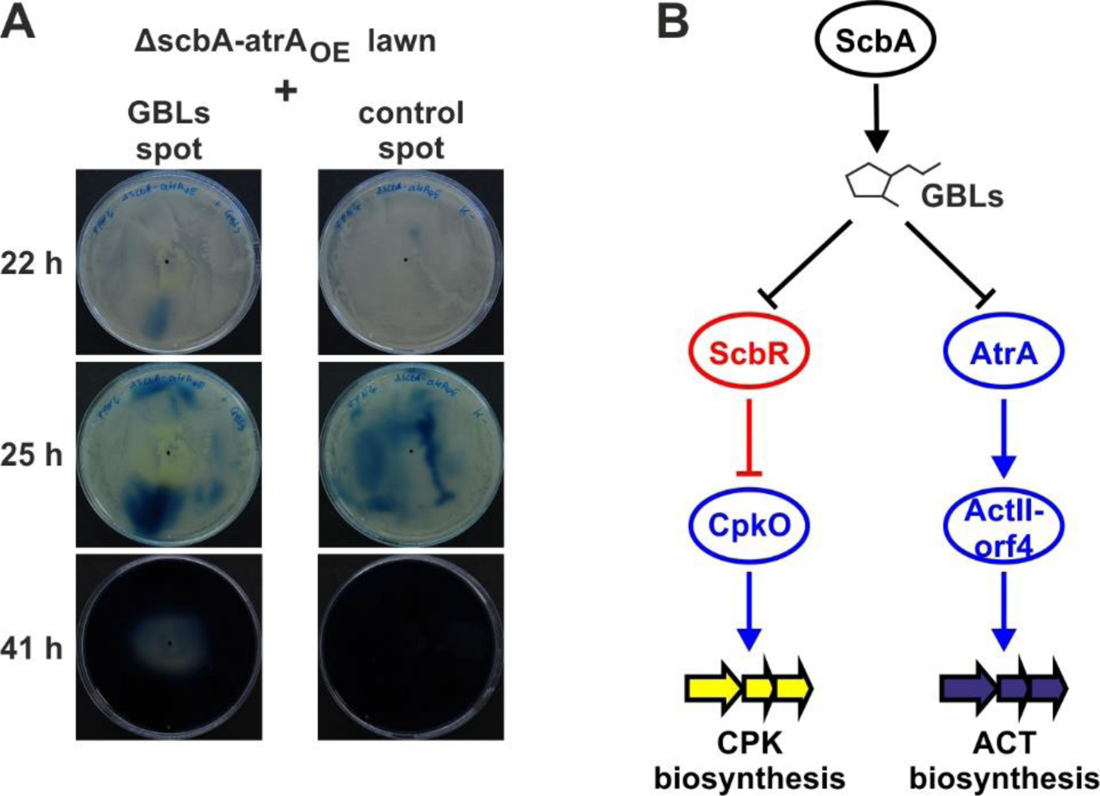
Simultaneous, GBL-mediated activation of coelimycin synthesis and inhibition of actinorhodin synthesis. A) Effect of GBL-rich extract on actinorhodin-overproducing *S. coelicolor* A3(2) Δ*scbA*-*atrA*OE, which is unable to produce CPK without exogenous GBLs. The negative control extract was obtained from the ΔscbA strain. B) Schematic representation of two different outcomes of GBL binding to the receptors ScbR and AtrA. The black arrow represents GBL synthesis. The black lines with bars represent the dissociation of GBL receptors from their target promoter regions upon GBL binding. Transcription activators and repressors are marked with blue and red, respectively, and their actions are marked with corresponding lines.

## DISCUSSION

The study of antibiotic production in *Streptomyces* has contributed greatly to further understanding of the underexplored field of quorum sensing in bacteria. The gamma-butyrolactone A-factor from *S. griseus* was indeed the first bacterial signalling molecule to be discovered in the 1960s (50). Among the identified GBL receptors are ArpA (regulator of streptomycin synthesis in *S. griseus*), JadR3 (jadomycin synthesis in *S. venezuelae*) (51), ScgR (natamycin synthesis in *S. chattanoogensis*) (52), FarA (synthesis of indigoidine, nucleoside antibiotics and D-cycloserine in *S. lavendulae* FRI-5) (53), BarA (virginiamycin production in *S. virginiae*) (54) and SscR (synthesis of unspecified secondary metabolites in *S. scabies* NBRC 12914) (55). Each of these proteins belongs to the TetR-like family of regulators (56). Two gamma-butyrolactone receptors are present in *Streptomyces coelicolor* A3(2): ScbR and the ScbR-like gamma-butyrolactone binding regulator SlbR. Although ScbR is a TetR-like protein encoded within the *cpk* biosynthetic gene cluster and has a well-established mechanism of action in the repression of coelimycin production, little is known about SlbR, which belongs to an unknown protein family. The pleiotropic effect of SlbR inhibition on ACT and RED production was shown without any specific mechanism (16). On the other hand, its binding within the *cpk* cluster (*scbR/A* promoter) suggests a specific pathway for CPK synthesis regulation that has never been shown to be relevant *in vivo* (18).

This work demonstrated that the well-characterized TetR-like protein AtrA is a gamma-butyrolactone receptor, as shown by *in vitro* GBL pull-down/mass spectrometry assay and EMSAs. The biological relevance of this effect is further supported by *in vivo* antibiotic production phenotype comparisons of several mutant strains (Δ*atrA*, *atrA*_OE_, Δ*scbA*φ and Δ*scbA*-*atrA*_OE_). These phenotypic differences, on the other hand, are explained by quantitative PCR analysis of the relevant transcripts (*atrA* and *actII-orf4* genes). Finally, the effect of gamma-butyrolactone-containing extract was demonstrated on the most potent ACT producer, Δ*scbA*-*atrA*_OE_, which at the same time lacks the ability to produce coelimycin. Externally added GBLs are able to inhibit this potent overproduction of actinorhodin and drive the synthesis of coelimycin instead. From these observations combined, it is evident how gamma-butyrolactones actually synchronize the life cycle progression of individual cells in a colony/culture not only by inducing coelimycin synthesis but at the same time, by actively blocking the synthesis of the later antibiotic actinorhodin (Fig. 6B). This is achieved by an elegant regulatory mechanism in which two TetR-like regulators, ScbR and AtrA, are able to receive the same signal (gamma-butyrolactones) but respond to it in the opposite manner. ScbR represses CPK synthesis SARP activator gene *cpkO* and dissociates from its promoter upon GBL-binding, leading to the production of yellow coelimycin. AtrA, on the other hand, is an activator of ACT synthesis SARP activator gene *actII-orf4* and dissociates from its promoter upon GBL-binding, causing the repression of actinorhodin synthesis. In this work, we described a comprehensive quorum sensing mechanism that ensures the synchronization of *S. coelicolor* A3(2) antibiotic production processes. This mechanism serves as a switch, ensuring that coelimycin and actinorhodin synthesis are mutually exclusive, timely-separated events. This is the scientific explanation for the simple observation of why in *S. coelicolor* A3(2) “there can be no actinorhodin while coelimycin is still being produced”.

Interestingly, MS analysis of GBL-containing extracts from the *ΔcpkO* and M1154 strains revealed the presence of three potentially new gamma-butyrolactones (tentatively named SCB9, SCB10 and SCB11), which were not reported by Sidda et al. (48). The amount of sample material was too low to elucidate the sample structure. In earlier work by Efremenkova and coworkers, six GBLs from *S. coelicolor* A3(2) were identified (see (57) for structures and references). Three of them (Acl-2a, Acl-2b and Acl-2d) are stereoisomers of SCB1 (58). Acl-2c has the same molecular mass as SCB1 and SCB2 but a different side chain structure. Acl-1a and Acl-1b are stereoisomers of SCB3. It cannot be excluded that putative SCB9, which has the same molecular mass as SCB1 and SCB2, corresponds to one of the Acl-2 series compounds. Similarly, putative SCB10 and SCB11, which have the same molecular mass as SCB3 and SCB7, may correspond to Acl-1a and Acl-1b.

A surprising finding of this study was the ability of ScbR2 to bind GBLs, as shown by the affinity capture method. This close homologue of ScbR is known as a “pseudo” GBL receptor, which shares some DNA targets with ScbR (59), but its DNA binding is not interrupted by the addition of GBL (60).

## SUPPLEMENTARY INFORMATION

**Additional file 1: Table S1.** Bacterial strains used in this work. **Table S2.** Plasmids and cosmids used in this work. **Table S3.** Oligonucleotides used in this work. **Figure S1.** HPLC-ESI-MS extracted ion chromatograms of GBLs from *S. coelicolor* cultures. Line colours represent calculated m/z values of the four ions ([M+Na]^+^, [M+H]^+^, [M-H2O+H]^+^, [M-2H2O+H]^+^) of GBLs (SCB1-SCB8) according to Sidda et al. (48), as indicated below chromatograms. Peaks are numbered with SCB numbers. Asterisks indicate putative new compounds SCB9, SCB10 and SCB11. **Figure S2.** HPLC-ESI-MS extracted ion chromatograms of GBLs from *S. coelicolor* M1154 cultures (initial GBLs) and from samples recovered from affinity capture experiment (protein names as indicated). Line colours represent calculated m/z values of the four ions ([M+Na]^+^, [M+H]^+^, [M-H2O+H]^+^, [M-2H2O+H]^+^) of GBLs (SCB1-SCB8) according to Sidda et al. (48), as indicated below chromatograms. Peaks are numbered with SCB numbers. Asterisks indicate putative new compounds SCB9, SCB10 and SCB11.

## ACKNOWLEDGEMENTS

This research was supported by Ludwik Hirszfeld Institute of Immunology and Experimental Therapy PAS.

## AUTHOR CONTRIBUTIONS

Conceptualization: BB; methodology: BB and MK; writing – original draft preparation: BB and MK; writing – review and editing: MK, BB, MW, KJP; performing experiments and formal analysis: BB, MK, MW, MD, AJ; resources: KJP; supervision: KJP. All the authors have read and approved the final manuscript.

## AVAILABILITY OF DATA AND MATERIALS

All the data generated and analysed in this study are included in this article and in the Additional files.

## DECLARATIONS

### Ethics approval and consent to participate

Not applicable.

### Consent for publication

All authors approved the manuscript.

### Competing interests

The authors declare no competing interests.

## Notes

### Competing Interest Statement

The authors have declared no competing interest.

