## Supplemental files for "The gamma-butyrolactone receptors ScbR and AtrA form a quorum sensing switch between coelimycin and actinorhodin synthesis in *Streptomyces coelicolor* A3(2)"

### SUPPLEMENTARY INFORMATION

Table S1. Bacterial strains used in this work

| Strain | Relevant genotype or description | Source or reference |
|---|---|---|
| ***Escherichia coli*** | | |
| DH5α | F$^-$ *endA1 glnV44 thi-1 recA1 relA1 gyrA96 deoR nupG* Φ80Δ*lacZ*ΔM15 Δ(*lacZYA-argF*)*U169*, *hsdR17*($r_K^-$ $m_K^+$), $\lambda^-$ | Promega |
| BL21(DE3)Star | F-, *ompT*, *hsdSB* (rB-, mB-), *gal*, *dcm*, *rne131* (DE3) | Novagen |
| BW25113/pIJ790 | *lacI*$^+$*rrnB*$_{T14}$ Δ*lacZWJ16 hsdR*514 Δ*araBAD*AH33 Δ*rhaBAD*$_{LD78}$ *rph-1* Δ*(araB–D)567* Δ*(rhaD–B)568* Δ*lacZ4787*(::*rrnB-3*) *hsdR514 rph-1* pIJ790<br><br>Recombineering strain harbouring arabinose-inducible RED genes on the plasmid pIJ790. | (1) |
| ET12567/pUZ8002 | strain for conjugal transfer of DNA from *E. coli* to *Streptomyces* (*dam dcm hsdS* Cam$^R$ Tet$^R$ on the bacterial chromosome; *tra* Kan$^R$ RP4 23 on pUZ8002) | (2) |
| ***Streptomyces coelicolor* A3(2)** | | |
| M145 | Wild type strain, *S. coelicolor* A3(2) (SCP1$^-$ SCP2$^-$) | (2) |
| M145-Φ (P132) | M145 with integrated empty pIJ10257 plasmid | This work |
| Δ*scbA* (M751) | M145 with in-frame deletion of *scbA* gene | Gift from Eriko Takano (3) |
| Δ*scbA*-Φ (P053) | M751 with integrated empty pIJ10257 plasmid | This work |
| M145-*atrA*$_{OE}$ (P330) | M145 with integrated pIJ10257-atrA$_{OE}$ | This work |
| Δ*scbA*-*atrA*$_{OE}$ (P331) | M751 with integrated pIJ10257-atrA$_{OE}$ | This work |
| Δ*atrA* (P332) | *atrA* gene replacement with apramycin resistance cassette *aac3(IV)* | This work |
| *atrA*$_{CO}$ (P333) | P332 with integrated pIJ10257-atrA$_{CO}$ | This work |
| Δ*atrA*-Φ (P334) | P332 with integrated empty pIJ10257 | This work |

Table S2. Plasmids and cosmids used in this work

| Name | Relevant genotype or description | Source or reference |
|---|---|---|
| pUC18 | Standard *E. coli* vector with a multiple cloning site (MCS) for DNA cloning | Thermo |
| pUC18-atrA | pUC18 with *atrA (SCO4118)* gene amplified with AtrA_F and AtrA_R primers, phosphorylated with T4 PNK (ThermoScientific) and cloned into SmaI site of the vector | This work |
| pET28a(+) | Novagen pET system overexpression plasmid | Novagen |
| pET28a-atrA | pET28a(+) containing *atrA* gene excised from pUC18-atrA with NdeI and HindIII and cloned into the same sites of the vector | This work |
| pUC18-slbR | pUC18 with *slbR (SCO0608)* gene amplified with SlbR_F and SlbR_R primers, phosphorylated with T4 PNK (ThermoScientific) and cloned into SmaI site of the vector | This work |
| pET28a-slbR | pET28a(+) containing *slbR* gene excised from pUC18-slbR with BamHI and HindIII and cloned into the same sites of the vector | This work |
| pIJ773 | Template to amplify *aac3(IV)* apramycin resistance cassette | (4) |
| StD72A | SuperCos1 cosmid carrying fragment of *S. coelicolor* A3(2) chromosome (bp 4516145 to 4549188) | http://strepdb.streptomyces.org.uk |
| StD72A-atrA$_{DM}$ | StD72A cosmid, in which *atrA* (*SCO4118*) gene sequence was replaced by means of PCR-targeting with an apramycin resistance gene *aac(3)IV* amplified using primers atrADMF and atrADMR1, and pIJ773 as a template | This work |
| pIJ10257 | ΦBT1 integrating overexpression plasmid containing strong constitutive promoter ermEp* | (5) |
| pIJ10257-atrA$_{CO}$ | pIJ10257 vector containing *atrA* gene with its promoter region amplified with primers atrAco_Uguru_R_HindIII and atrAco_Uguru_F_KpnI and introduced into KpnI and HindIII digested vector. The complemented sequence is the same as in (6) | This work, (6) |
| pIJ10257-atrA$_{OE}$ | pIJ10257 vector containing *atrA* gene excised from pUC18-atrA with NdeI and HindIII and cloned into the same sites of the vector | This work |
| pTZ57R/T | T-vector from InstT/A Cloning kit for direct cloning of PCR products | Thermo |

| | | |
|---|---|---|
| pTZ57R-pactII-orf4 | pTZ57R/T containing promoter fragment of *actII-orf4* | (7) |
| pFLUXH | ΦBT1 integrating reporter plasmid with a promoterless luciferase operon *luxCDAEB* and hygromycin resistance cassette | (8) |
| pFLUXH-patrA | pFLUXH containing promoter fragment of *atrA* gene amplified with atrA-6 and atrA-11 and cloned by Gibson assembly into BamHI and NdeI-digested vector | This work |

Table S3. Oligonucleotides used in this work. Restriction sites are in bold. The homology arms for Gibson assembly are underlined.

| Name | Sequence 5'-3' | Restriction sites | Application/amplified fragment |
|---|---|---|---|
| AtrA_F | **CATATG**CATGTTCAGGATTCTCATTGG | NdeI | *atrA* gene for cloning in expression plasmid |
| AtrA_R | **AAGCTT**TCACACCGGCCGCGACCGC | HindIII | |
| atrADMF | TCTCCGGGGGGAGACGTCATTACCGGGGGATTGTCTATGATTCCGGGGATCCGTCGACC | | *aac3(IV)* apramycin resistance cassette for *atrA* gene replacement |
| atrADMR1 | GAAGGAGATACGGGCCCCCGACGACGTCGCCGCGTCTCATGTAGGCTGGAGCTGCTTC | | |
| atrA-5 | ATATCGGCATCGATGCATGATCACGCAGGTCAGCGTGGG | | *atrA* promoter region (fragment *patrA*) for cloning into pFLUXH |
| atrA-11 | ATATCGGCATCGATGCATGCTAGACGACCGTGATCTCGGCC | | |
| atrAX | GCTCAACTATGTTCTTTCCCACGGTG | | Verification of *atrA* deletion |
| atrAY | TCACTCGTCTGTGCGGACTTCACG | | |
| SlbR_F | **GGATCC**CATATGTCCGAGAGCACGATGCAGTCCG | BamHI | Amplification of *slbR* |
| SlbR_R | **AAGCTT**TCAGCTGCCGTTCCGCGCG | HindIII | |
| pTZBAM700 | IRDye700-ATGCAGGCCTCTGCA | | amplification and IRDye700 labelling of fragments for EMSA cloned in pTZ57R/T |
| pTZXBA700 | IRDye700-TCGGTACCTCGCGAA | | |
| EMSAFLUXF-FAM | 6FAM-CCAGTACTTCGCGAAAGC | | amplification and IRDye700 labelling of fragments for EMSA cloned in pFLUXH |
| EMSAFLUXR-FAM | 6FAM-ATGATGAACGAGATCTTCTTCG | | |

Figure S1 HPLC-ESI-MS extracted ion chromatograms of GBLs from *S. coelicolor* cultures. Line colours represent calculated m/z values of the four ions ($[M+Na]^+$, $[M+H]^+$, $[M-H2O+H]^+$, $[M-2H2O+H]^+$) of GBLs (SCB1-SCB8) according to Sidda et al. (48), as indicated below chromatograms. Peaks are numbered with SCB numbers. Asterisks indicate putative new compounds SCB9, SCB10 and SCB11.

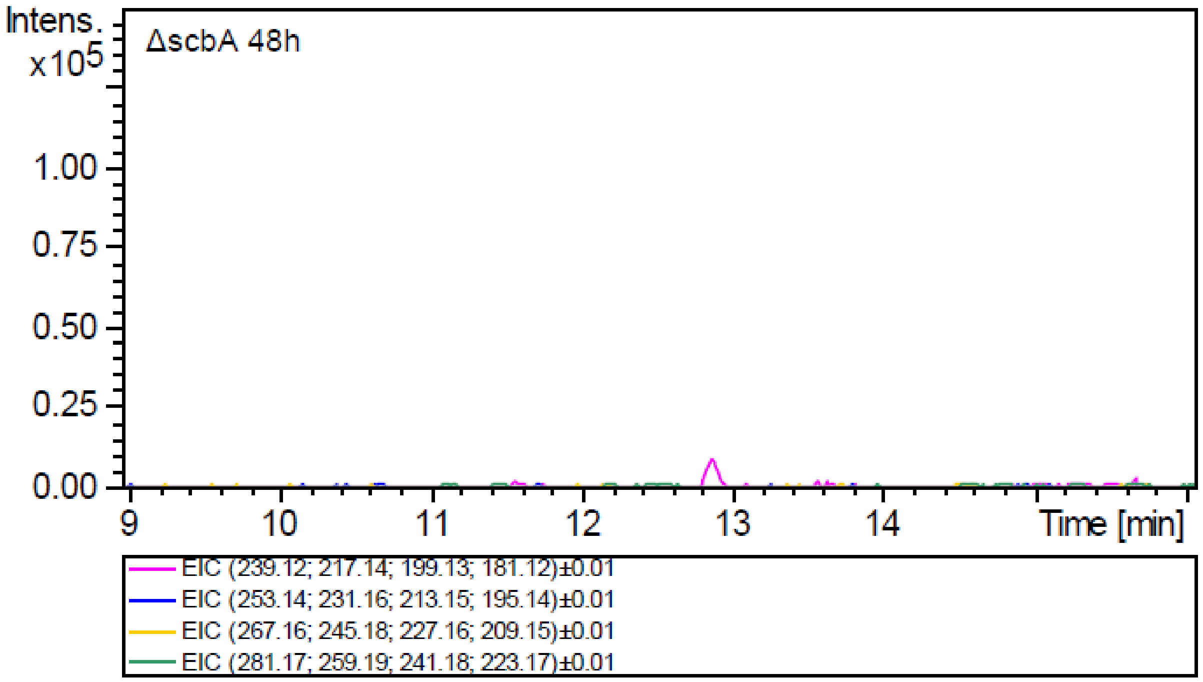

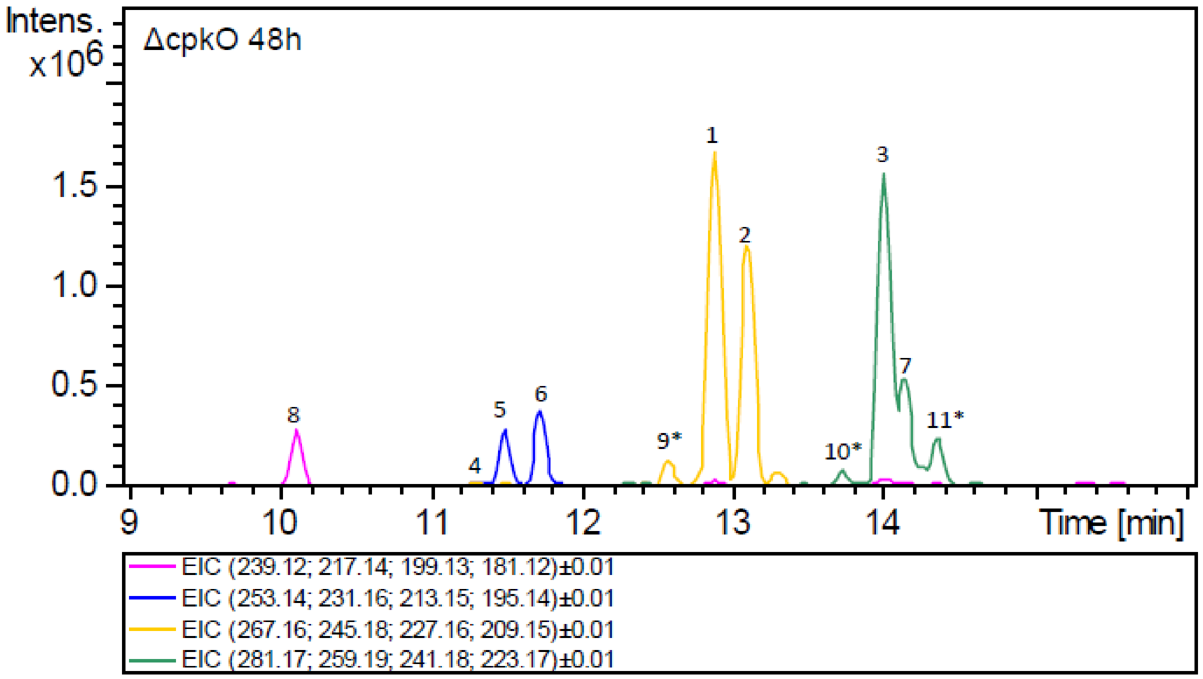

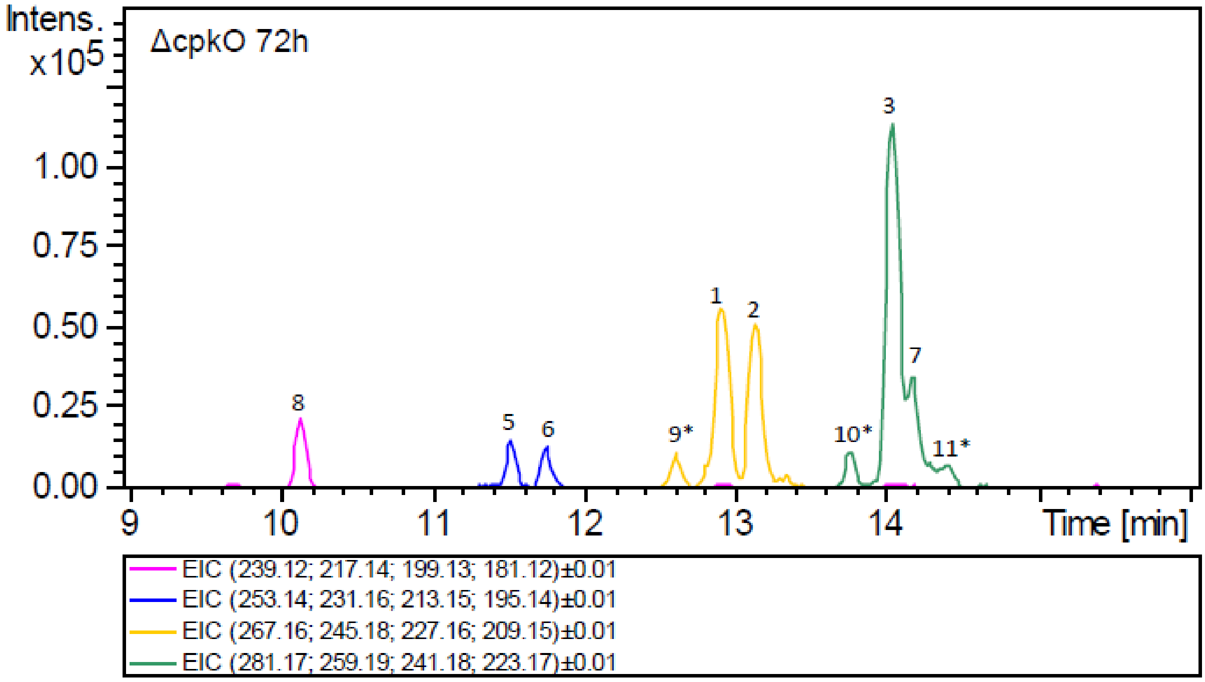
Intens.
x10^5
ΔcpkO 72h
1.00
0.75
0.50
0.25
0.00
8
5
6
9*
1
2
10*
3
7
11*
9
10
11
12
13
14
Time [min]
EIC (239.12; 217.14; 199.13; 181.12)±0.01
EIC (253.14; 231.16; 213.15; 195.14)±0.01
EIC (267.16; 245.18; 227.16; 209.15)±0.01
EIC (281.17; 259.19; 241.18; 223.17)±0.01

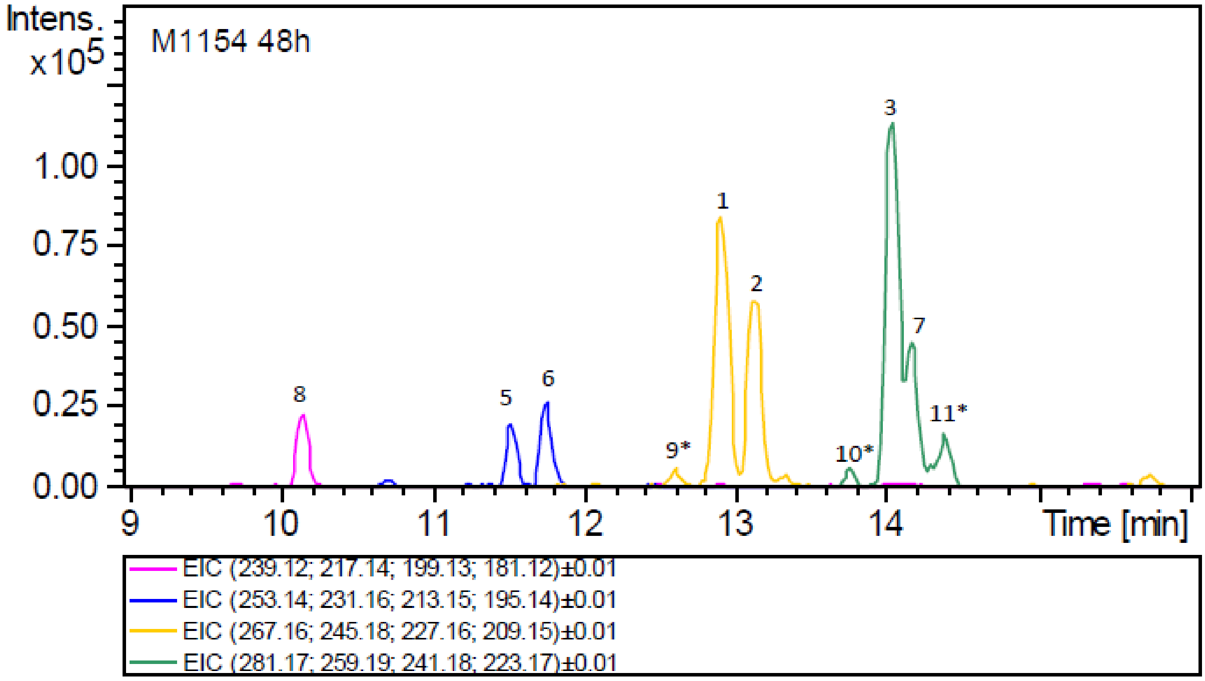
Intens.
x10^5
M1154 48h
1.00
0.75
0.50
0.25
0.00
8
5
6
9*
1
2
10*
3
7
11*
9
10
11
12
13
14
Time [min]
EIC (239.12; 217.14; 199.13; 181.12)±0.01
EIC (253.14; 231.16; 213.15; 195.14)±0.01
EIC (267.16; 245.18; 227.16; 209.15)±0.01
EIC (281.17; 259.19; 241.18; 223.17)±0.01

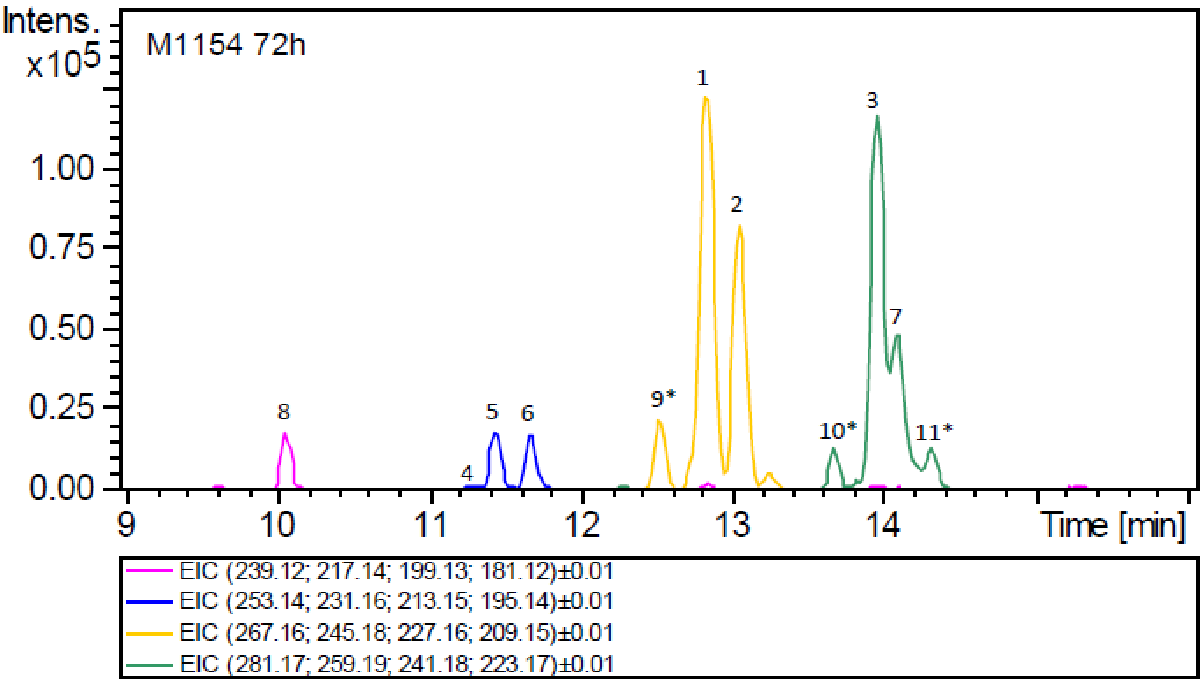
Intens.
x10^5
M1154 72h
1.00
0.75
0.50
0.25
0.00
1
2
3
4
5
6
7
8
9*
10*
11*
9
10
11
12
13
14
Time [min]
EIC (239.12; 217.14; 199.13; 181.12)±0.01
EIC (253.14; 231.16; 213.15; 195.14)±0.01
EIC (267.16; 245.18; 227.16; 209.15)±0.01
EIC (281.17; 259.19; 241.18; 223.17)±0.01

Figure S2. HPLC-ESI-MS extracted ion chromatograms of GBLs from *S. coelicolor* M1154 cultures (initial GBLs) and from samples recovered from affinity capture experiment (protein names as indicated). Line colours represent calculated m/z values of the four ions ($[M+Na]^+$, $[M+H]^+$, $[M-H2O+H]^+$, $[M-2H2O+H]^+$) of GBLs (SCB1-SCB8) according to Sidda et al. (48), as indicated below chromatograms. Peaks are numbered with SCB numbers. Asterisks indicate putative new compounds SCB9, SCB10 and SCB11.

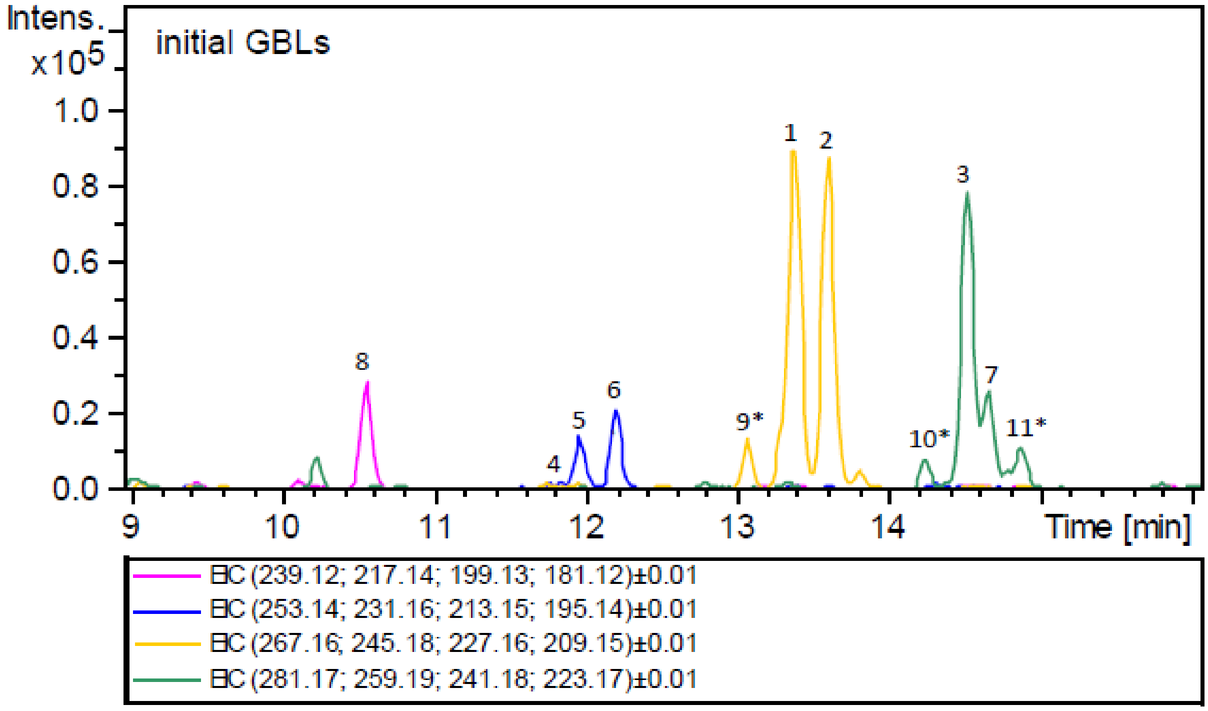

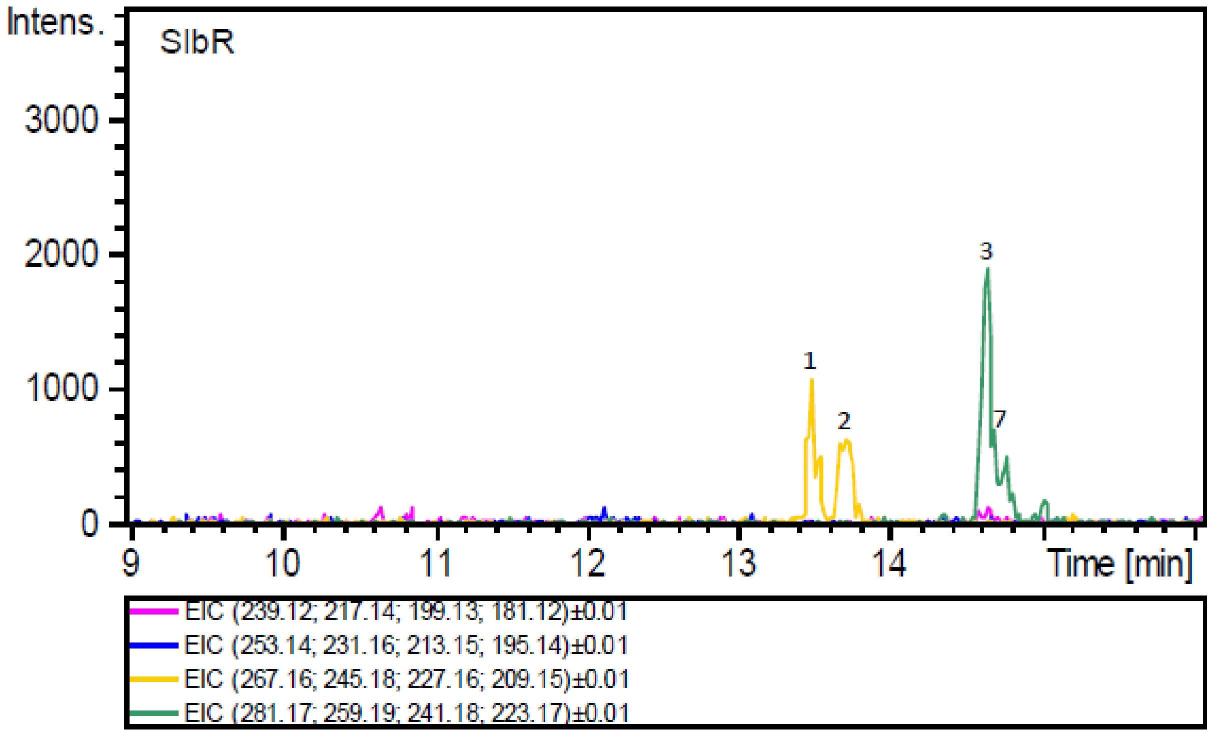

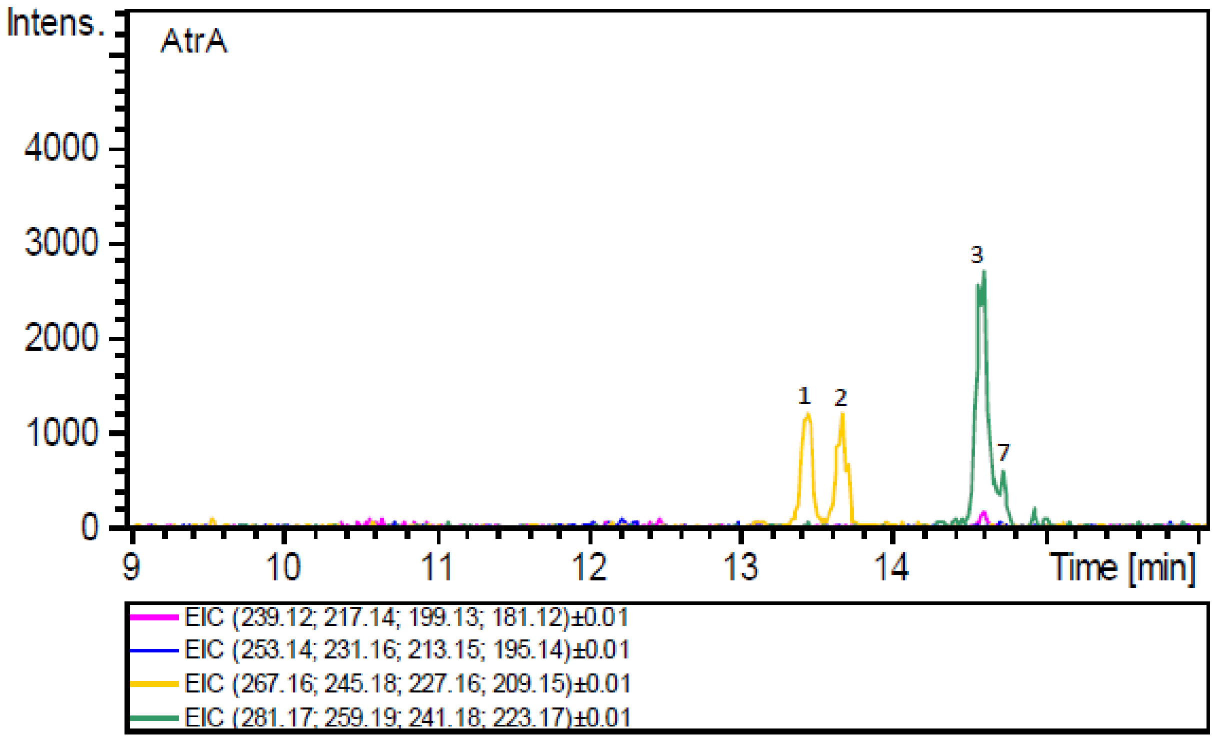
AtrA
Intens.
4000
3000
2000
1000
0
9
10
11
12
13
14
Time [min]
1
2
3
7
EIC (239.12; 217.14; 199.13; 181.12)±0.01
EIC (253.14; 231.16; 213.15; 195.14)±0.01
EIC (267.16; 245.18; 227.16; 209.15)±0.01
EIC (281.17; 259.19; 241.18; 223.17)±0.01

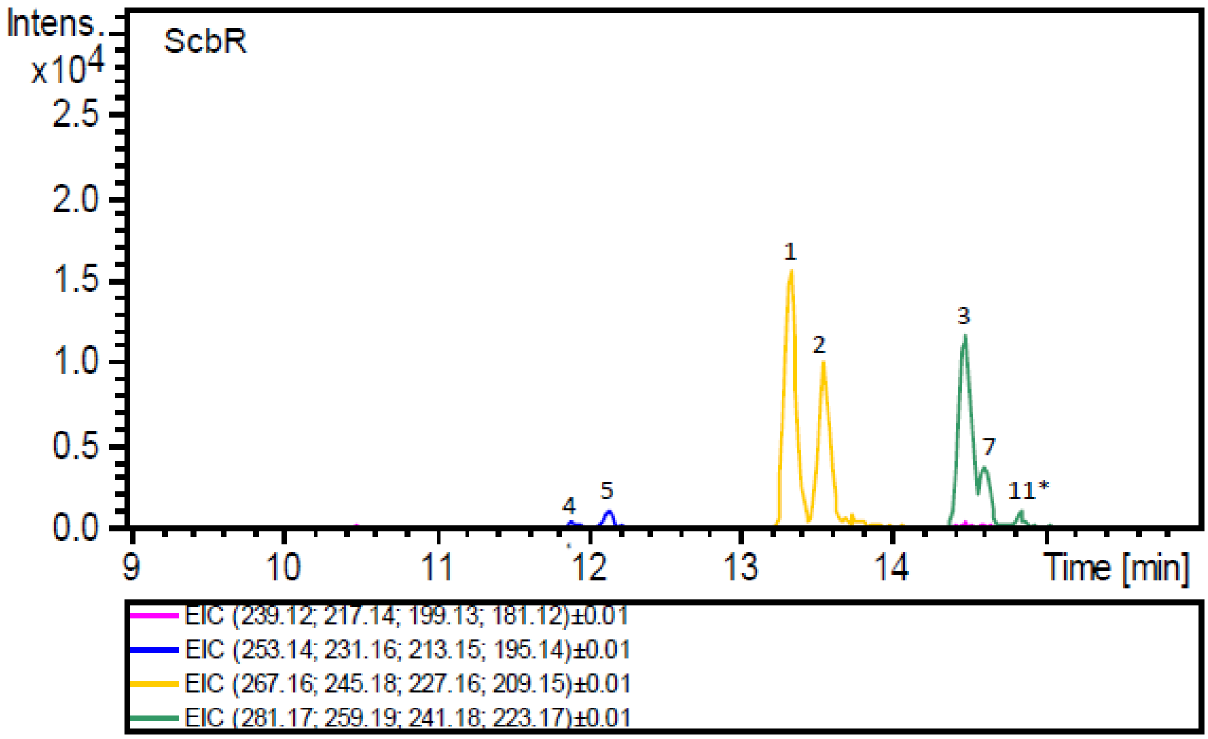
ScbR
Intens.
x10^4
2.5
2.0
1.5
1.0
0.5
0.0
9
10
11
12
13
14
Time [min]
1
2
3
4
5
7
11*
EIC (239.12; 217.14; 199.13; 181.12)±0.01
EIC (253.14; 231.16; 213.15; 195.14)±0.01
EIC (267.16; 245.18; 227.16; 209.15)±0.01
EIC (281.17; 259.19; 241.18; 223.17)±0.01

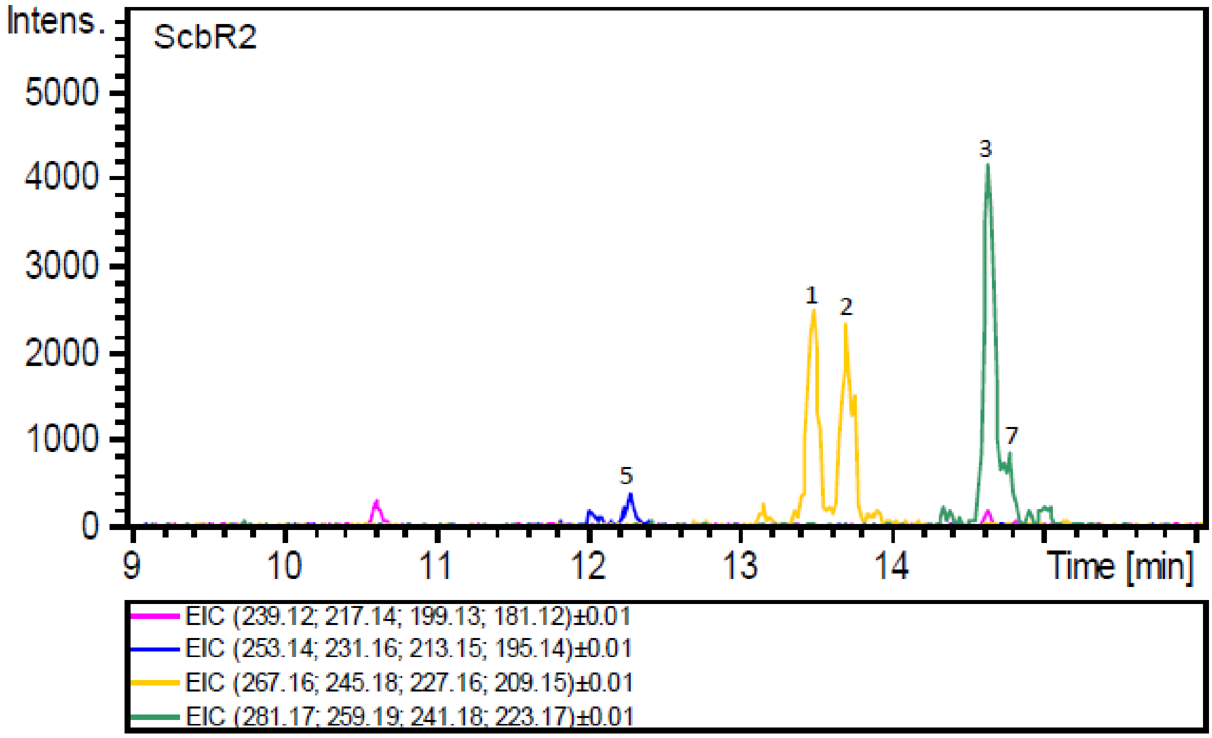

Intens.
ScbR2
5000
4000
3000
2000
1000
0
1
2
3
5
7
9
10
11
12
13
14
Time [min]
EIC (239.12; 217.14; 199.13; 181.12)±0.01
EIC (253.14; 231.16; 213.15; 195.14)±0.01
EIC (267.16; 245.18; 227.16; 209.15)±0.01
EIC (281.17; 259.19; 241.18; 223.17)±0.01

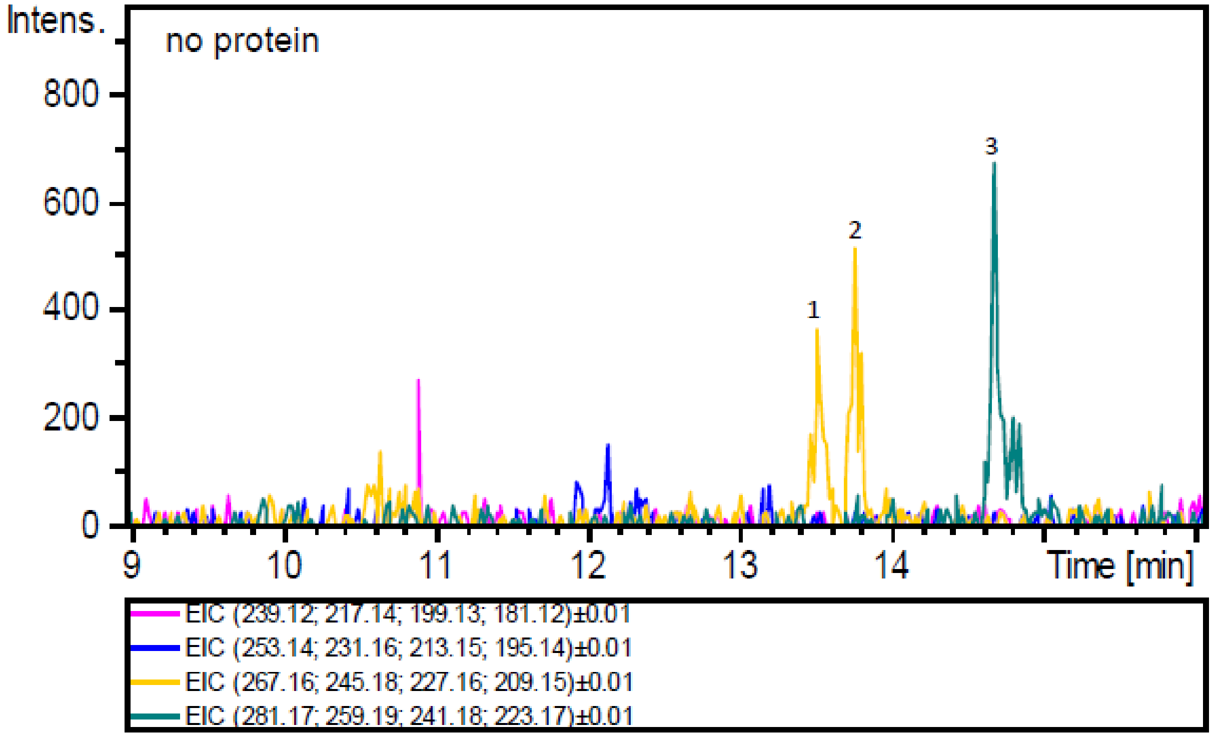

Intens.
no protein
800
600
400
200
0
1
2
3
9
10
11
12
13
14
Time [min]
EIC (239.12; 217.14; 199.13; 181.12)±0.01
EIC (253.14; 231.16; 213.15; 195.14)±0.01
EIC (267.16; 245.18; 227.16; 209.15)±0.01
EIC (281.17; 259.19; 241.18; 223.17)±0.01

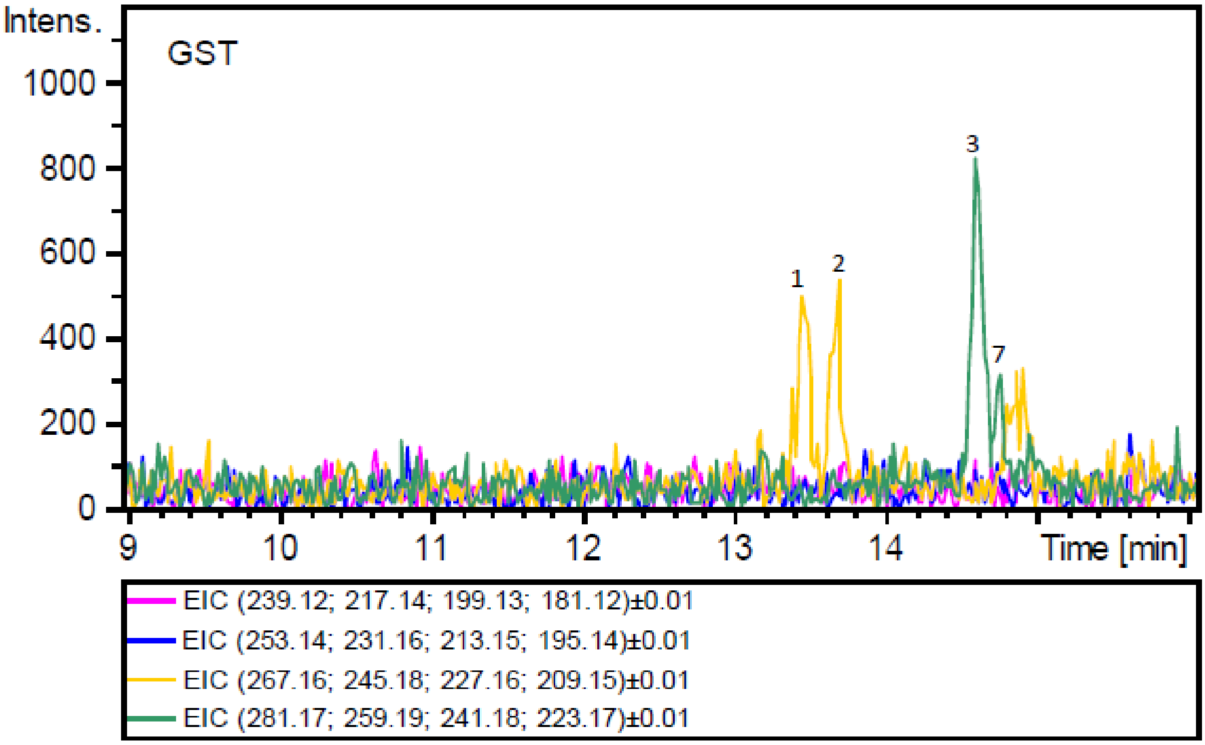

#### SUPPLEMENTARY INFORMATION REFERENCES
